# Identification of novel drug-drug interactions: towards precision medicine in women’s reproductive health

**DOI:** 10.1101/2024.11.25.624843

**Authors:** Pablo Garcia-Acero, Ismael Henarejos-Castillo, Patricia Sebastian-Leon, Antonio Parraga-Leo, Juan Antonio Garcia-Velasco, Patricia Diaz-Gimeno

**Author notes:** Corresponding author: Patricia Diaz-Gimeno., IVI Foundation, Instituto de Investigación Sanitaria La Fe, Av. Fernando Abril Martorell 106. Torre A, Planta 1ª, 46026 Valencia, Spain. Ismael Henarejos-Castillo and Pablo Garcia-Acero are joint first authors.

## Abstract

Drug-drug interactions (DDIs) may occur when two or more drugs are taken together, leading to undesired side effects or potential synergistic effects. Most clinical effects of drug combinations have not been assessed in clinical trials. Therefore, predicting DDIs can provide better patient management, avoid drug combinations that can negatively affect patient care, and exploit potential synergistic combinations to improve current therapies in women healthcare. For this purpose, a DDI prediction model was built to describe relevant drug combinations affecting reproductive treatments. Approved drug features (chemical structure of drugs, side effects, targets, enzymes, carriers and transporters, pathways, protein-protein interactions, and interaction profile fingerprints) were obtained. A unified predictive score revealed unknown DDIs between reproductive and commonly used drugs and their associated clinical effects on reproductive health. The performance of the prediction model was validated using known DDIs.

This prediction model accurately predicted known interactions (AUROC = 0.9876) and identified 2,991 new DDIs between 192 drugs used in different female reproductive conditions and other drugs used to treat unrelated conditions. These DDIs included 836 between drugs used for in-vitro fertilization. Most new DDIs involved estradiol, acetaminophen, bupivacaine, risperidone, and follitropin. Follitropin, bupivacaine, and gonadorelin had the highest discovery rate (42%, 32%, and 25% respectively). Some were expected to improve current therapies (n=23), while others would cause harmful effects (n=11). We also predicted twelve DDIs between oral contraceptives and HIV drugs that could compromise their efficacy. Overall, these results show the importance of DDIs studies to personalize female reproductive therapies.

## 1. Introduction

Drug-drug interactions (DDIs) may occur when two or more drugs are co-administered to a patient. In some cases, the drugs may act synergistically to amplify the pharmacological effect, but in others, the interactions can reduce pharmacodynamic efficacy, cause adverse drug events (ADEs) [1], or variable effectiveness [2]. DDIs are emerging as a public health concern, with a recent study estimating that DDIs account for 5% of all hospital admissions [2].

Considering the vast number of over-the-counter and prescription drugs on the market, the average American adult takes three drugs per day [3], and 22.4% of adults were dispensed with five or more drugs [4], with women more likely to take polypharmacy than men [5]. In the context of female reproductive medicine, the DDIs between contraceptives, antiretroviral drugs [i.e., for human immunodeficiency virus (HIV)], and other unrelated drugs, are alarming clinicians [6,7]. Although there are several ongoing studies related to DDIs on contraception [8–10], patients undergoing fertility treatments sometimes need to take multiple drugs to treat their complex conditions, causing potential DDIs and ADEs. Indeed, in a recent study of 440 patients with polycystic ovary syndrome (PCOS), up to nine different contraceptives and infertility drugs were co-administered, causing 26.1% of these patients to present DDIs, which required close monitoring [11]. With the uncertainty of the long-term effects of coronavirus disease 2019 (COVID-19) on women’s reproductive health [12,13], and the combination of drugs used to manage its systemic symptoms, patients with COVID-19 may similarly be at risk. Taken together, predicting DDIs could offer substantial benefits, especially for women undergoing assisted reproduction treatments (ARTs). After further validation, the identification of DDIs may allow clinicians to offer safer, more effective therapeutic strategies, while opening paths to develop new combination of treatments [14].

Further, studying DDIs through *in-silico* approaches based on predictive models is both cost-effective and efficient, compared to conventional *in-vitro* or *in-vivo* approaches. Most *in-silico* approaches to predict new DDIs are based on the similarities between drugs (e.g., mainly chemical similarity) [15], but also their comparable side effects [16,17], known interactions with other drugs, denoted as interaction profile fingerprint (IPF) [18], shared targets, enzymes, carriers, transporters, or molecular signaling pathways [19,20], or proximity of targets in the human interactome [21]. Indeed, Vilar *et al.* published a detailed methodology for integrating a reference database of known DDIs, with all the drug similarities, to significantly improve their predictions [18]. To our knowledge, drugs used in any capacity in women’s reproductive health, hereafter referred as women’s reproductive health drugs (WRHDs), have not been assessed in-depth by these methods. Herein, we present a framework to analyze known DDIs and predict new DDIs in the context of women’s reproductive health. Thus, the novel WRHDs interactions described here cover different reproductive statuses (from preconception to menopause); associated reproductive diseases or conditions (including uterine, ovarian, and menstrual disorders); clinical application (*in-vitro* fertilization [IVF], and ovarian stimulation); relevant diseases (HIV, COVID-19) contraceptives; and drugs not specifically used for reproduction, but potentially taken concurrently (e.g., anesthetics, related to surgical-procedures, or to treat conditions outside of reproductive health). For the first time in gynecology, this study innovatively compiled information of drugs used to treat women’s reproductive diseases/conditions, with their known DDIs (extracted from pharmacological databases), to predict novel DDIs, that can help future research for clinical decision-making in gynecology, and ultimately, achieve safer and more effective treatments in women’s reproductive medicine.

## 2. Material and methods

### 2.1. Compilation of approved drugs indicated for women’s reproductive medicine

Drug data related to women’s reproductive statuses or clinical application [i.e., preconception, infertility, menopause, IVF], diseases and conditions (i.e., menstrual, uterine, ovarian, or other reproductive disorders) were classified according to the query terms listed in **Supplemental Table 1A**. Guidelines of the European Society of Human Reproduction and Embryology (ESHRE) were consulted to identify approved drugs indicated for the management of endometriosis [22], premature ovarian insufficiency (POI) [23], recurrent pregnancy loss (RPL) [24], polycystic ovary syndrome (PCOS) [25], Turner syndrome [26], ovarian stimulation [27], and oocyte retrieval procedures [28], until September 2020. To complete the list of approved WRHDs, we then consulted PubMed, the DrugBank database [29], and ClinicalTrials.gov (for drugs in phase IV clinical trials, used in women with reproductive diseases between 2013 to 2020).

### 2.2. Building the model to predict drug-drug interactions involving women’s reproductive health drugs

Known DDIs (n = 117,002) for each WRHD included in the study were obtained from DrugBank [29]. Six drug features were evaluated: (i) chemical structure (based on the premise that if drug A and drug B interact to produce a biological effect, then drugs with chemical similarity to drug A or drug B can produce the same effect when they are combined) [15] obtained from DrugBank; (ii) drug targets, enzymes, transporters, and carriers obtained from DrugBank (since DDIs can occur when molecular entities are shared) [20]; (iii) ADEs reported in the Side Effect Resource (SIDER) [30]; (iv) targeted biological pathways included in Kyoto Encyclopedia of Genes and Genomes (KEGG) [31]; (v) proximity of drug targets in the human interactome [32,33] from the CCSB Interactome Database; (vi) the IPF of each drug (which assumes that if two drugs share similar interaction partners with other drugs, then these two drugs could also interact between themselves) [18]. Afterwards, the methodological protocol reported by Vilar *et al.* 2014 to build a prediction model was strictly followed [18]. Briefly, we built a reference matrix (M1) with the DDIs for each WRHD, followed by matrices for each of the six features (M2), where we calculated drug similarities with the Tanimoto index [34]. Crossing M1 by each M2 matrix formed another six prediction matrices (M3) containing the predicted scores. These scores were then integrated into a single score for each DDI, through a principal component analysis (PCA), using the ROCR package in R [35]. Regarding establishing a threshold of confidence for the novel interactions predicted, Vilar et al., recommend setting to the third quartile of the distribution of predicted scores. We calculated our third quartile from the distribution of scores from an internal validation set. Briefly, we used the ROCR package to calculate the AUROC (10-fold cross validation) when predicting known interactions from DrugBank [16,18]. After filtering the predicted DDIs by this threshold of confidence, the potential biological effect of each DDI was classified according to annotations in DrugBank, assessing their potential clinical impact. Next, we performed a descriptive study of these WRHDs DDIs encompassing IVF drugs. Finally, we investigated possible interactions with COVID-19 and HIV drugs, and interactions of IVF drugs with non-gynecological drugs.

## 3. Results

### 3.1. Classifying drugs used in women’s reproductive health

Based on our searches of the ESHRE guidelines, ClinicalTrials.gov, DrugBank, and PubMed, we identified 192 WRHDs associated with 51 different reproductive conditions or diseases (**Supplemental Table 1B**), classified into the following categories: preconception (47 drugs), infertility and IVF (58 drugs), menopause (9 drugs), uterine diseases (83 drugs), ovarian diseases (67 drugs), menstrual disorders (17 drugs), and other reproductive disorders (73 drugs).

### 3.2. Known and novel drug-drug interactions involving women’s reproductive health drugs

Including the 192 unique WRHDs, a total of 4,014 approved drugs were retrieved from DrugBank. An exhaustive search in DrugBank yielded 117,002 known DDIs among the 192 WRHDs and the other 3,822 approved drugs (**Supplemental Table 2**), which we then used to validate model’s performance (AUROC = 0.9876 ± 0.0149). To highlight novel interactions, we set the threshold at the third quartile (0.7418) of the validation set. After filtering by this threshold, we obtained 2,991 novel predicted interactions (**Supplemental Table 3**) between the WRHDs and the drugs for non-gynecological indications, a novel discovery of 2.5% with respect to the 117,002 known DDIs used in the validation set. Notably, 15 of the 192 WRHDs (7.8%) did not have any known or predicted interactions, and eight of these were IVF drugs [i.e., luteinizing hormone, lutropin alfa, urofollitropin, menotropins, human chorionic gonadotropin (hCG) and its alpha subunit (hCGα), cetrorelix, and ganirelix]. Our findings highlight that DDIs can not only affect a variety of conditions and diseases, (including uterine disorders, PCOS, and infertility), but also produce ADEs and changes in the pharmacokinetics and pharmacodynamics of WRHDs. Estradiol was distinguished as the WRHD with the most interactions from the preconception and the menopause groups; acetaminophen for menstrual disorders; bupivacaine in uterus diseases; risperidone for other reproductive diseases; and follitropin (a recombinant form of follicle stimulating hormone) for infertility and ovarian disease groups (**Figure 1A**). On the other hand, follitropin, bupivacaine, and gonadorelin were the WRHDs with the highest DDI discovery rate (**Figure 1B**).

**Figure 1.**
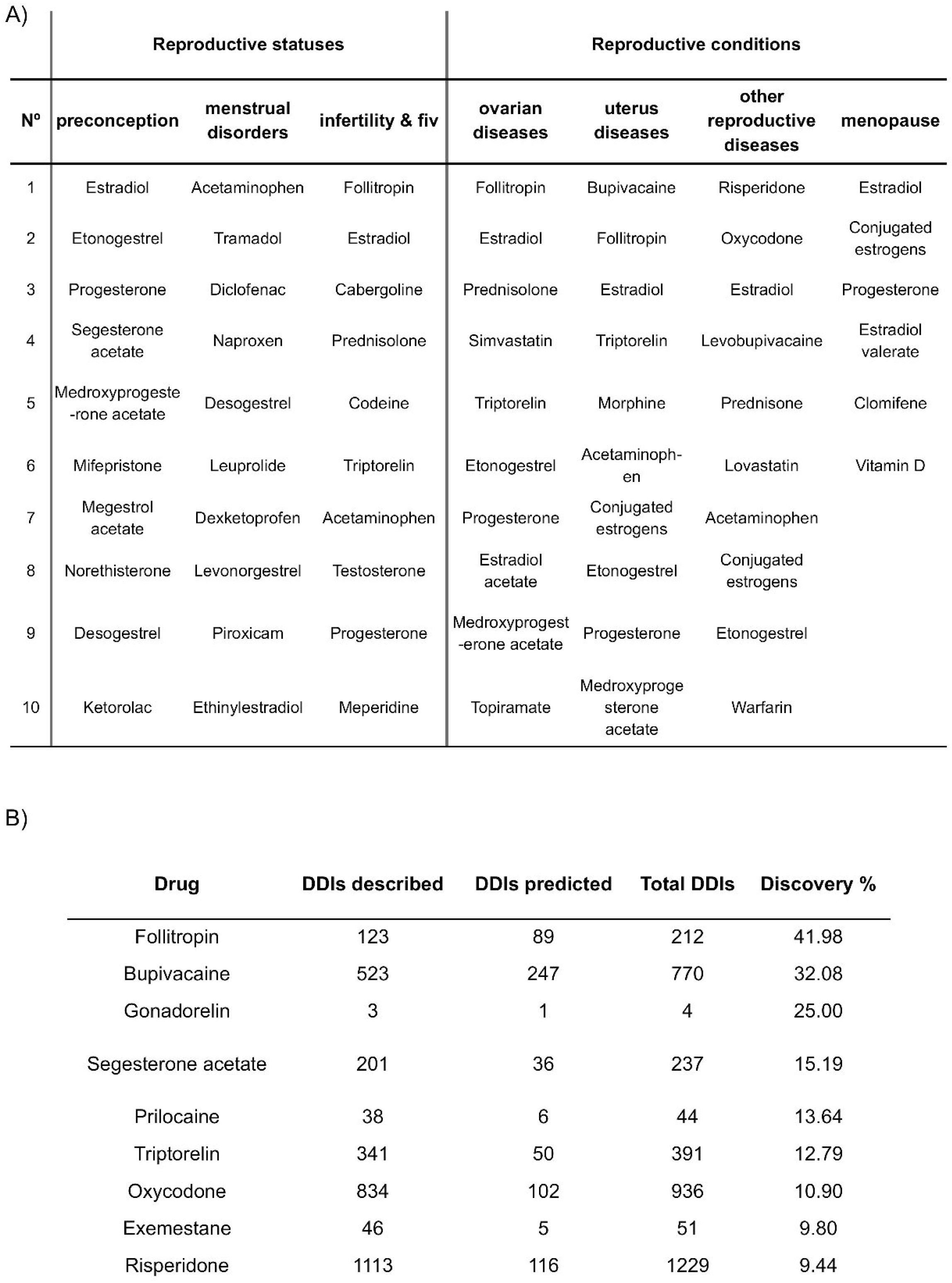
Discovery rate of drug-drug interactions for female reproductive drugs. The Discovery rate represents the proportion between predicted drug-drug interactions and already described interactions. The top 10 female reproductive drugs by highest rate are represented in the figure. The number of described interactions is shown in blue, whereas the number of predicted, novel interactions are shown in yellow. Follitropin, gonadorelin, and triptorelin came on top, with 41.98%, 25%, and 12.79% novel discovery rates, respectively.

Regarding the interactions between WRHDs and contraceptives or HIV drugs, we predicted twelve novel DDIs for amphotericin B, and three for alitretinoin. We found that three COVID-19-related drugs were under investigation to treat reproductive disorders (i.e., azithromycin for preeclampsia, ibuprofen for preeclampsia and endometriosis, and dexamethasone for PCOS). Further, we predicted that heparin (indicated for RPL) may improve the therapeutic efficacy of chloroquine and methylprednisolone (two drugs used to treat COVID-19); four novel interactions between azithromycin and antihypertensive agents [producing an extended interval between the heart contracting and relaxing (QT prolongation)]; nine between Ibuprofen and drugs indicated for gastrointestinal disorders, hypertension, diabetes, asthma, mental disorders, and inflammation; as well as six between dexamethasone and antihypertensives, mental disorder drugs, and antineoplastics (**Supplemental Table 4**).

### 3.3. Drug-drug interactions predicted to pose a problem on women’s reproductive health

Analyzing the predictions from our model, we report novel DDIs for our WRHDs with relevant clinical impact (**Table 1**). Patients co-administered fentanyl and follitropin may have an increased risk or severity of cardiac arrhythmia, while cyclosporine may decrease the efficacy of follitropin. In concordance with previous studies, we found that combining progesterone and estradiol might increase the risk or severity of liver damage. Further, we also found that prednisolone might accelerate the metabolism of midazolam and lidocaine, which are both used for anesthesia during oocyte retrieval procedures. Finally, we highlight an interesting prediction in which patients taking cabergoline may experience reduced estradiol metabolism during controlled ovarian stimulation.

**Table 1.**
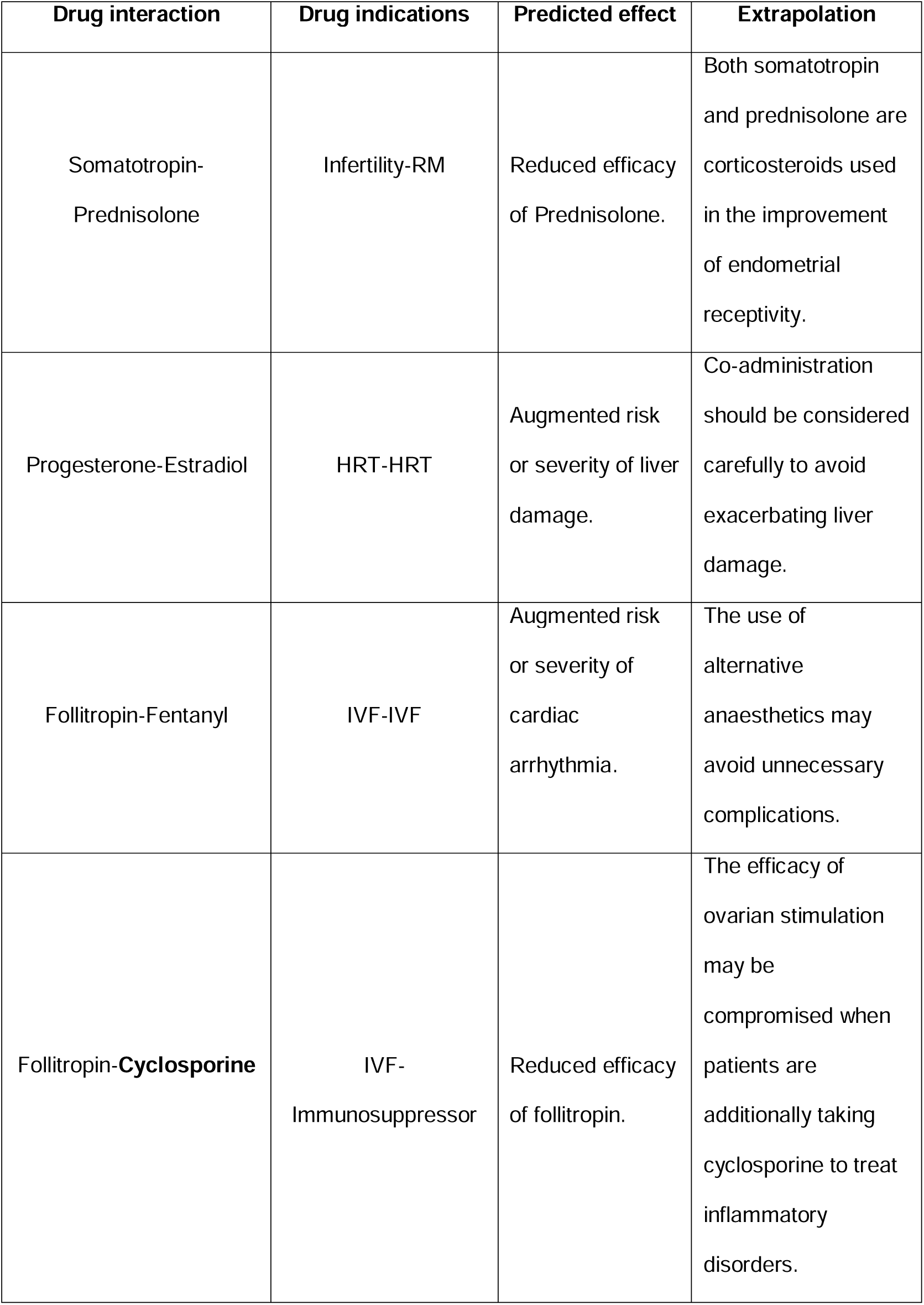
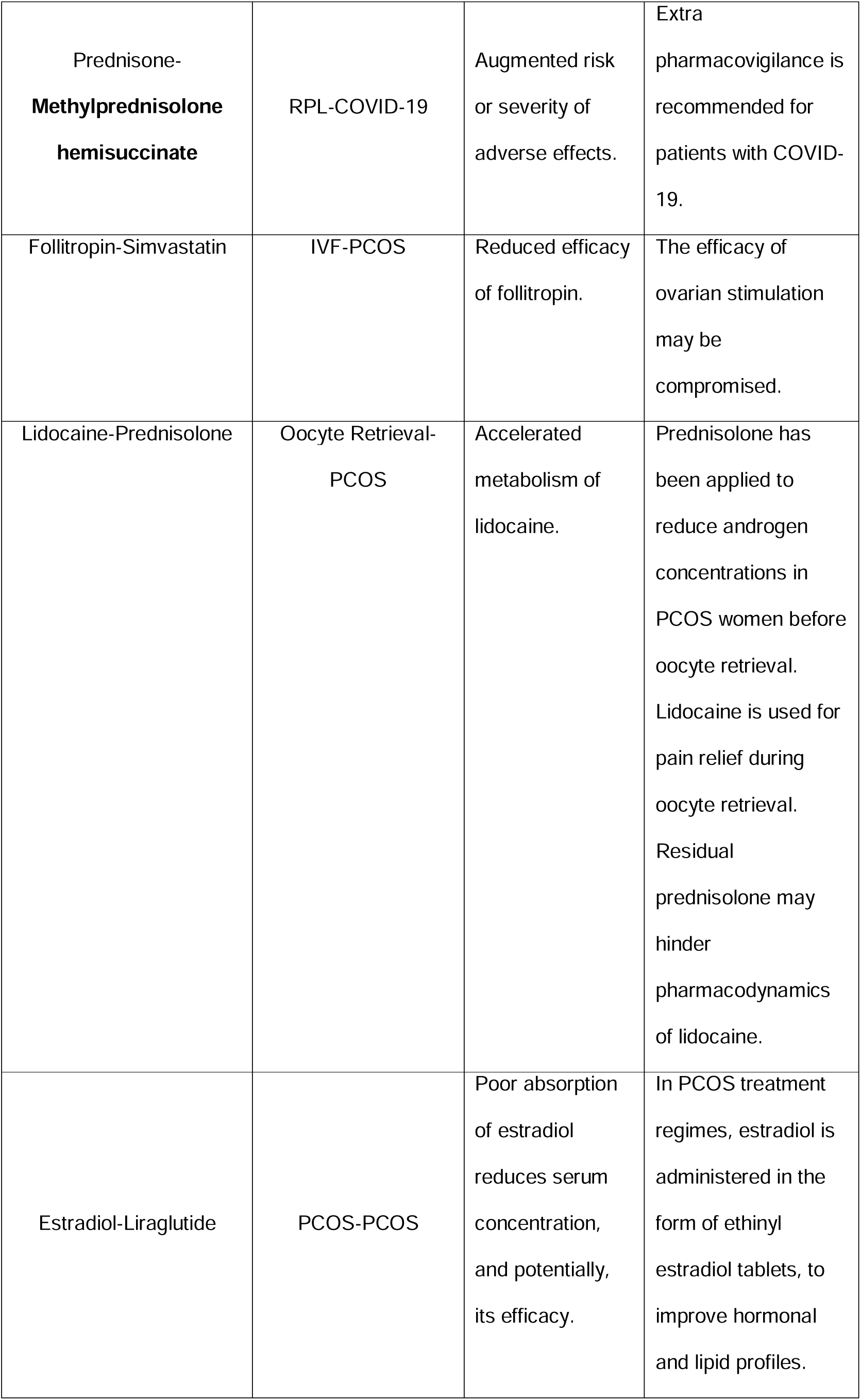
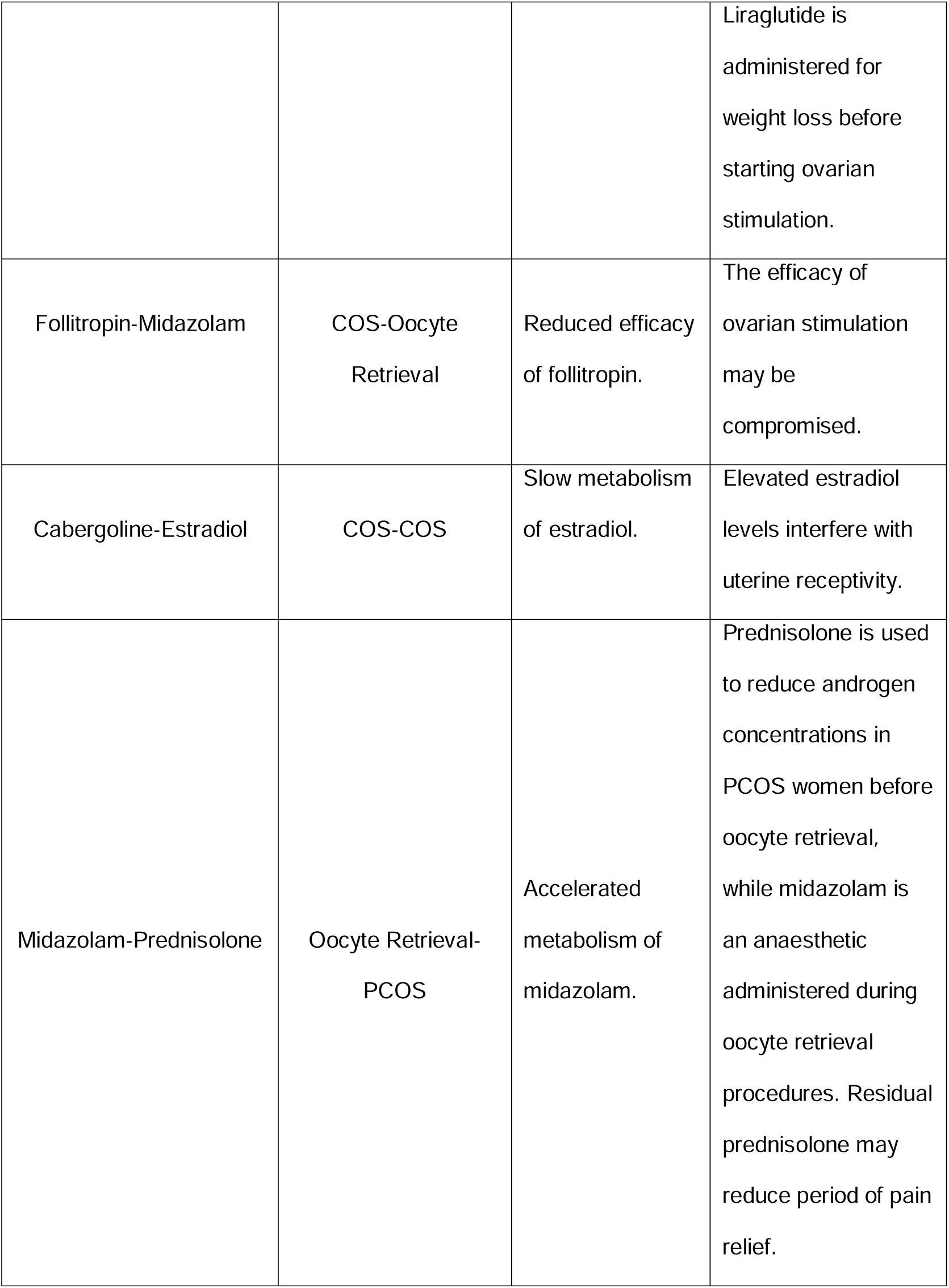
Interactions predicted to pose clinical conflicts. Eleven previously unknown drug-drug interactions that pose potential clinical conflicts. Drugs currently used for non-gynecological indications are highlighted in bold. ART, assisted reproduction technology; COS, controlled ovarian stimulation; HRT, hormone replacement therapy; IVF, in-vitro fertilization; PCOS, polycystic ovary syndrome; RM, recurrent miscarriage; RPL, recurrent pregnancy loss.

### 3.4. Drug-drug interactions predicted to enhance the therapeutic efficacy of current women’s reproductive health

We identified 23 DDIs that could potentially improve the therapeutic efficacy of one drug when combined with another (**Table 2**). Specifically, we predicted that when vitamin D and estradiol are co-administered (e.g., in the treatment of PCOS), the accelerated metabolism of estradiol may occur. Interestingly, four of the nine drug combinations predicted to improve the efficacy of triptorelin (used for patients with PCOS) involved other drugs commonly administered to patients with PCOS [i.e., isotretinoin (indicated for PCOS-related acne), levocarnitine (L-carnitine; indicated for insulin resistance- or obesity-related PCOS), folic acid, and pyridoxine (also indicated for endometriosis)]. We also note interactions between isotretinoin and gonadorelin, and methyldopa (indicated for preeclampsia) and chlorothiazide (currently indicated for hypertension).

**Table 2.**
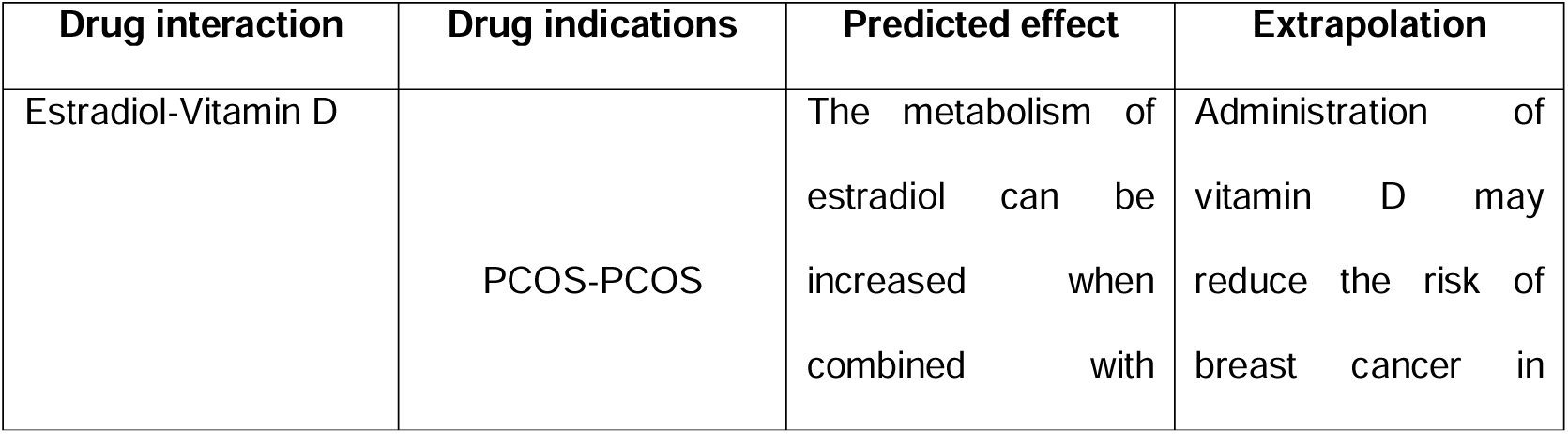
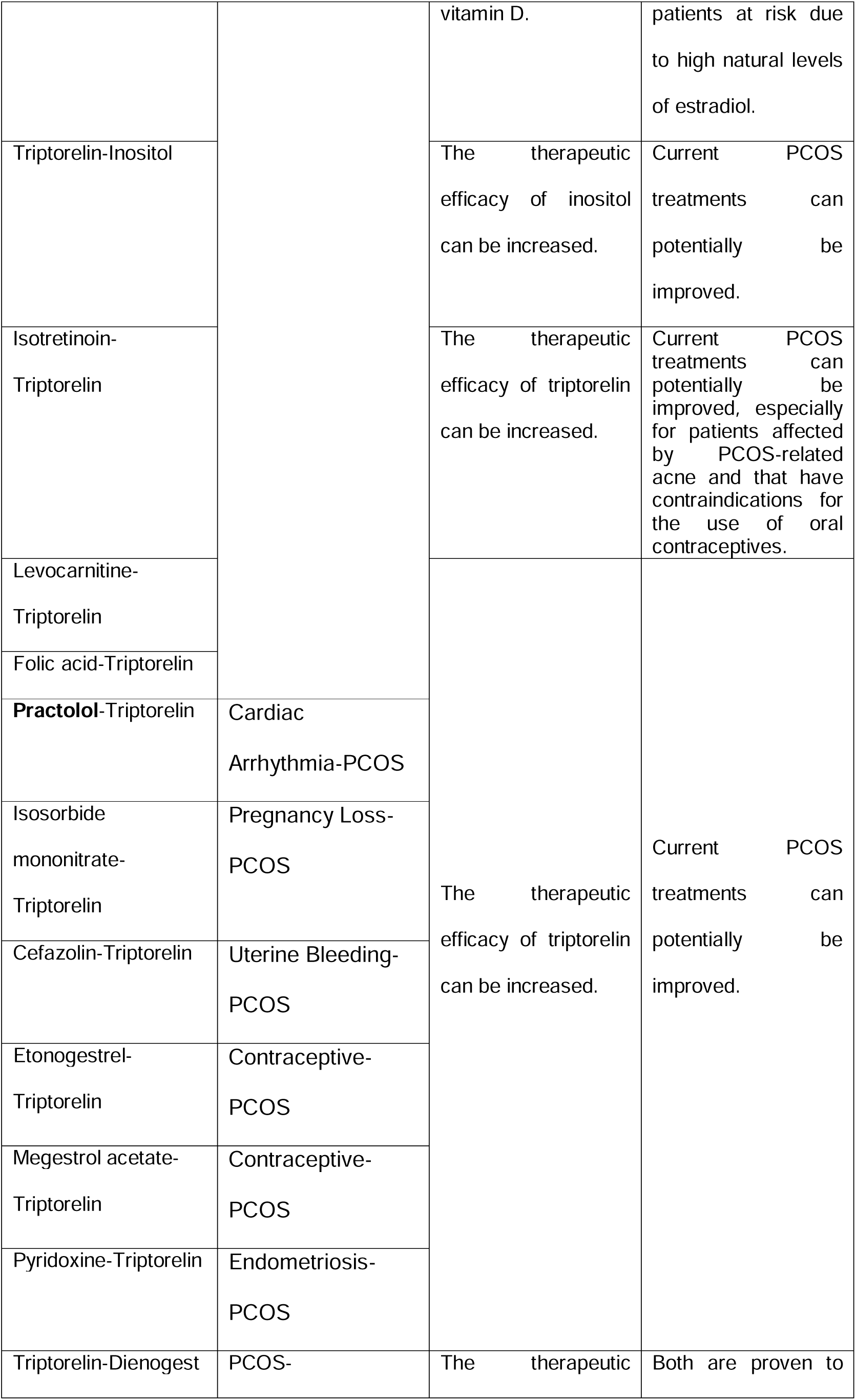
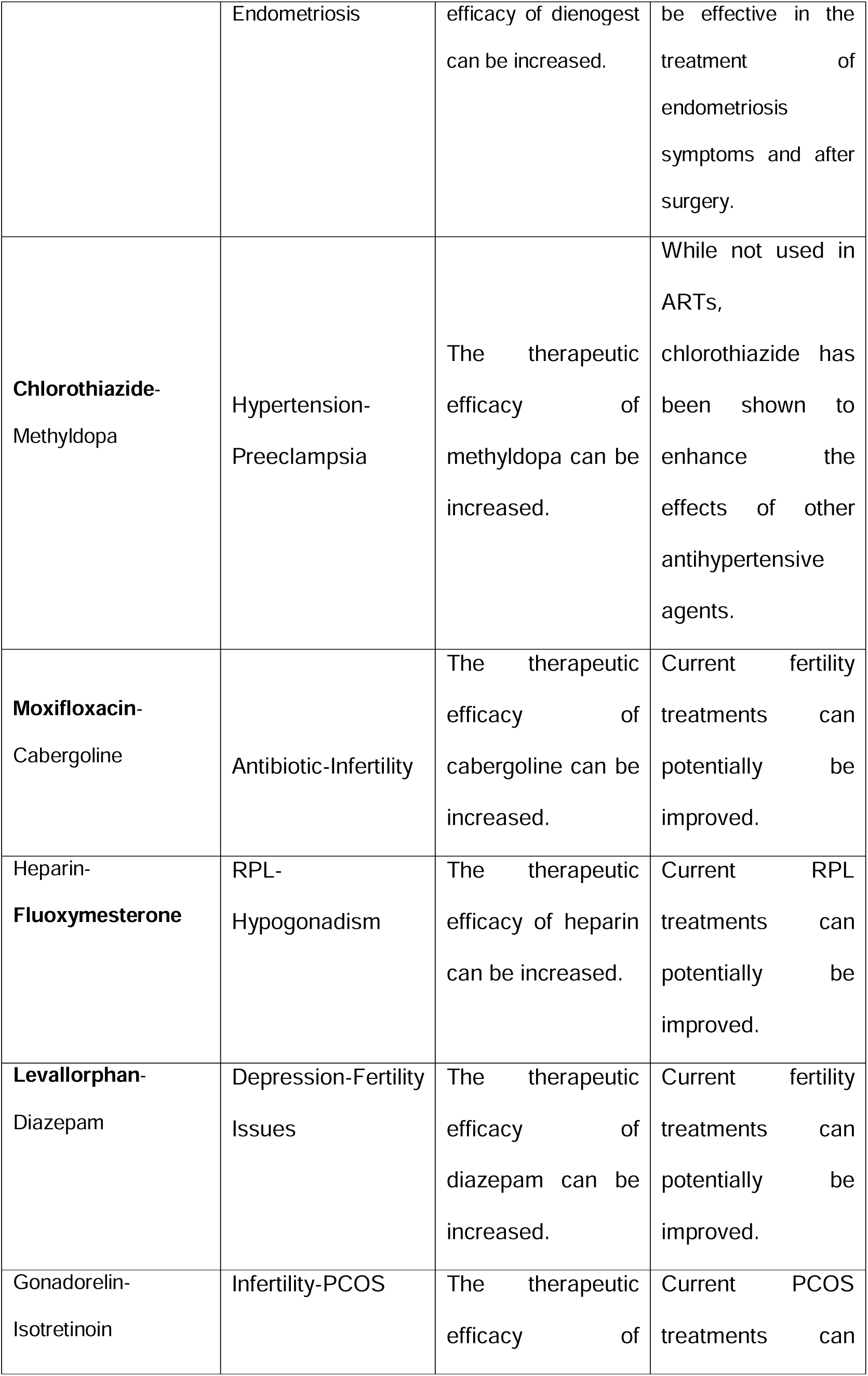
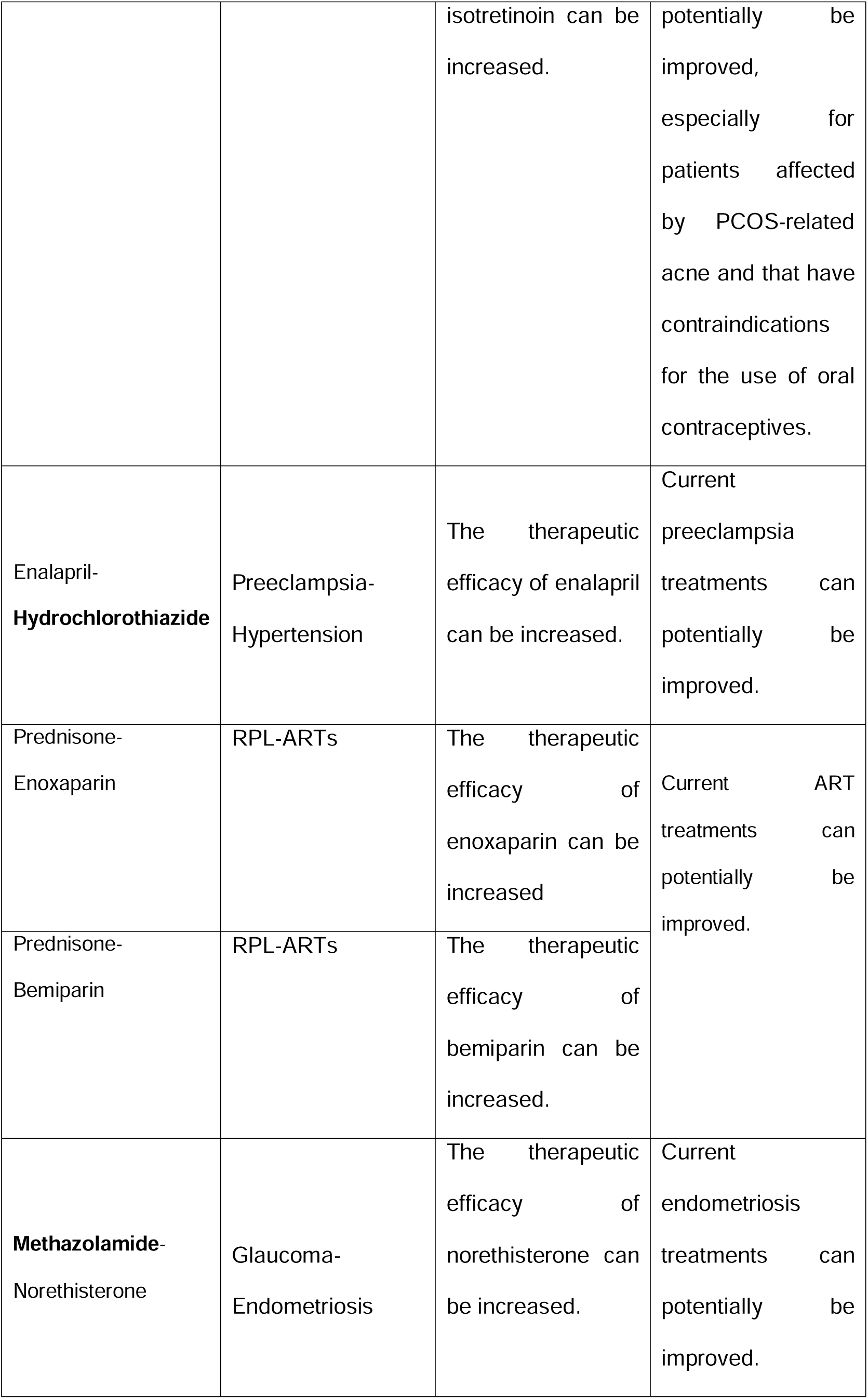
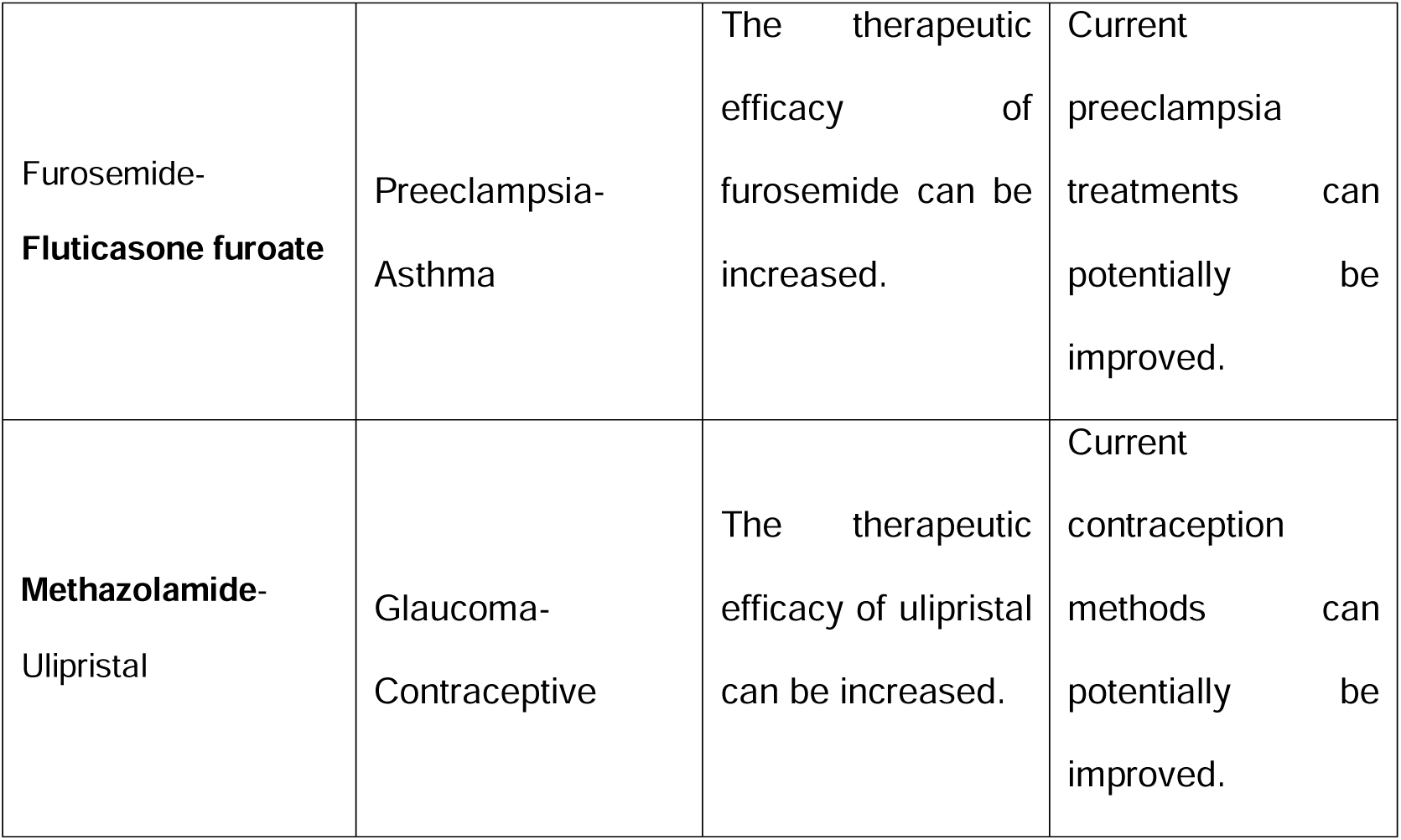
Interactions predicted to enhance pharmacological efficacy. Twenty-three previously unknown interactions that enhance the pharmacological efficacy, of women’s reproductive health drugs. Drugs currently used for non-gynecological indications are highlighted in bold. ART, assisted reproduction technology; HRT, hormone replacement therapy; PCOS, polycystic ovary syndrome; RPL, recurrent pregnancy loss.

### 3.5. Predicted conflicts between IVF drugs and drugs for non-gynecological indications

The subset of IVF drugs (n=58) (listed in **Supplemental Table 5**) from the WRHDs was used to analyze the predicted interactions with the most common drugs used in the general clinical setting. We predicted 836 novel interactions between IVF drugs and drugs approved for non-gynecological indications [mainly for asthma, allergies, anti-inflammation, analgesics, sedatives, and diabetes (**Figure 2A and Supplemental Table 5**). Specifically, follitropin was the IVF drug with the most predicted DDIs (n=82). Estradiol mainly interacted with drugs related to asthma and allergy (n=9), or diabetes (n=3), while cabergoline mainly interacted with anti-inflammatories, analgesics, and sedatives (n=21). Networks showcasing the distribution of normalized interactions between WRHDs or IVF drugs and their therapeutic indications are shown in **Figure 2B** and **2C**, respectively. Estradiol had the most interactions between IVF drugs (n=7), while bupivacaine had the most among all WRHDs (n=21). Drugs indicated for uterine disorders showed the highest number of DDIs with the rest of the WRHDs (n=48), whereas drugs used for infertility had the largest number of DDIs with IVF drugs (n=20). Specific data for these interactions is shown in **Supplemental Table 5**.

**Figure 2.**
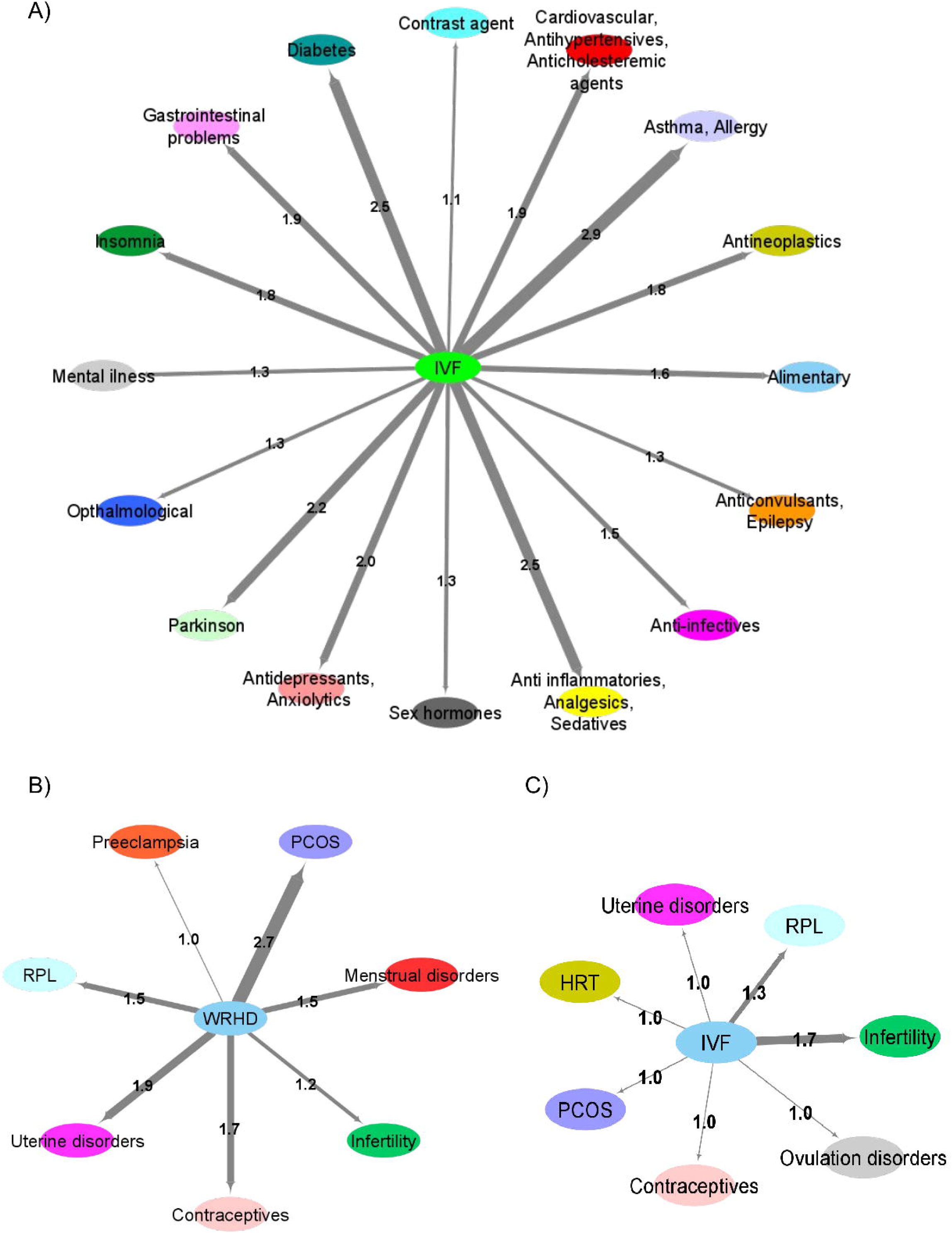
Interactions between drug categories: Circular nodes represent a set of drugs related to different indications while edges are the number of drug interactions predicted between both sets of drugs (the thickness of the edges are proportional to the number of drug-drug interactions between different set of drugs) **(A)** Predicted drug-drug interactions between female reproductive drugs and other approved drugs considering the indications. **(B)** Predicted drug-drug interactions between female reproductive drugs and IVF drugs considering the indications **(C)** Predicted drug-drug interactions between IVF drugs considering the indications.

## 4. Discussion

To our knowledge, this is the first study to use a predictive model for DDIs in women’s reproductive health, which could potentially improve the clinical management of ART treatments. In addition to summarizing known interactions at a systematic level, we adapted a robust prediction model to discover unknown DDIs that can compromise the efficacy and safety of current ART treatments, or, alternatively, improve their effectiveness [14]. Individually analyzing the DDIs allowed us to elucidate their potential clinical implication and advise clinicians to observe the reduced efficacy, unknown ADEs, or therapeutic benefits, resulting from the combination of drugs used in female reproductive medicine, and set the stage for clinical trials that aim to improve the treatment of women’s diseases/conditions.

The predictive model was built using the biological and pharmacological data of 4,014 approved drugs retrieved from DrugBank. This data encompassed chemical structures, known ADEs, shared targets, carriers, enzymes, transporters, pathways, interactome data, and interaction profiles [15–20], following the methodology proposed by Vilar et al., who were pioneers in this type of studies [18]. The capacity of our model to accurately predict the 117,002 known interactions for the 4,014 drugs from DrugBank demonstrates the reliability of the novel interactions we predicted in this study.

### 4.1. Identifying clinically relevant drug-drug interactions to aid ART practitioners

Our model highlighted follitropin (indicated for ovarian stimulation) at the forefront of the novel DDIs discovery. However, considering the normalized proportion of predicted interactions with drugs indicated for non-gynecological disorders, we found that IVF drugs interact mainly with asthma and allergy drugs, followed closely by drugs for diabetes (both with estradiol coming on top), and anti-inflammatories, analgesics, and sedatives drugs (in the case of cabergoline). Taken together, our findings reinforced that female ART drugs can interact with others indicated for non-gynecological disorders.

Upon close examination of the predicted interactions, here we report different recommendations for drug combinations according to the effect of the predicted interaction. We predicted nine positive DDIs that can increase the efficacy of triptorelin suggested for treatment of endometriosis [36,37], which is often co-administered with oral contraceptives We also found a potential synergistic effect for the combination of isotretinoin and gonadorelin [38,39], which are recommended for the treatment of PCOS-related acne when oral contraceptives are contraindicated. According to our results, the interaction between these two drugs would enhance the efficacy of isotretinoin. We also predicted a synergistic interaction between triptorelin and levocarnitine. Recent studies suggest that levocarnitine improves pregnancy rates and alleviates symptoms of ovarian dysfunction [40,41]. Therefore, the addition of triptorelin to levocarnitine-based treatment regimens merits prospective clinical validation in patients with endometriosis. Further, we predicted that vitamin D promotes the metabolism of estradiol [42,43]. Women with unbalanced levels of endogenous estradiol are at risk of developing endometriosis [44]. Therefore, a prospective validation of vitamin D for the treatment of endometriosis warrants attention. Finally, we predicted a positive interaction of methyldopa (an antihypertensive agent used for preeclampsia) and chlorothiazide (currently indicated for hypertension, but not ARTs) [45]. Interestingly, chlorothiazide has enhanced the effectiveness of other antihypertensive drugs [29], but its combination with methyldopa has not yet been evaluated clinically, especially in the context of preeclampsia.

Alternatively, based on the possible adverse DDIs we predicted, we warn that the combination of fentanyl (used as an anesthetic during oocyte retrieval procedures) [46] and follitropin (used for ovarian stimulation) [47] may increase the risk or severity of cardiac arrhythmia. Although the administration of these two drugs is not concurrent *per se*, a slower metabolism of follitropin may overlap with fentanyl administration. We also validate that cabergoline slows the metabolism of estradiol, supporting the findings by Lin *et al.* [48], which argued that the addition of cabergoline to GnRH antagonist protocols may be detrimental to uterine receptivity by maintaining high serum estradiol levels, thus supporting the strategy of embryo cryopreservation following GnRH antagonist protocol [49,50]. Further, despite the widespread use of estradiol and progesterone in female reproductive medicine [51,52], the risk of liver damage when administering both drugs has been previously described (possibly through elevated levels of liver aminotransferases) [53,54] and of developing cholestasis or other hepatic conditions, even if sometimes this risk is known and accepted [55–58]. We predicted the potential increase in this risk, reinforcing the validity of our drug-drug interactions predictions. Regarding treatments for reproductive conditions, most interactions predicted as harmful were found between drugs indicated for PCOS.

### 4.2. Clinical Implications

This study highlights the advantages of using predictive models to report unknown DDIs, advance post-market pharmacovigilance by forewarning clinicians of unreported ADEs and/or potential reduced clinical effectiveness, providing insight on alternative drug combinations that may improve the effectiveness of currently available treatments, and ultimately, patient’s fertility.

### 4.3. Research implications

This is the first study that analyzes and compiles together a list of known DDIs for women’s reproductive health, and predicts compromising and synergistic DDIs, not only increasing awareness of DDIs in the context of women’s reproductive health but also identifying research gaps that can be addressed by future clinical studies aimed to develop novel treatments for female patients undergoing ARTs.

### 4.4. Strengths and limitations

Between 2009 and 2017, the US Food and Drug Administration (FDA) approved 302 new drugs [59], but their post-market pharmacovigilance was limited with respect to evaluating possible interactions between ART drugs and other previously approved drugs in clinical use. In this regard, the pharmaceutical industry, regulatory agencies, public health services, and patients, would benefit from the development of robust prediction models to discover novel DDIs. However, the model is limited to what is stored in the databases consulted, with many drugs lacking information about their targets, ADEs, and more; the incompleteness of the human interactome; and the assumption that the databases are well curated, among other limitations. Furthermore, prospective studies are needed to validate the DDIs predicted by our model.

Nonetheless, this study sheds light on DDIs relevant to women’s reproductive health, revealing previously unknown DDIs that could compromise the efficacy of ART treatments, or boost the therapeutic effects of drugs such as follitropin or triptorelin. We acknowledge that further experimental and clinical evidence is needed to validate our predictions, however, we note that our prediction model was highly sensitive when evaluating currently known DDIs, making it a promising tool for precision medicine. Indeed, the information generated from this study could be implemented in institutional computerized prescription systems that alert clinicians of potential conflicts now when the medication is ordered for clinical observational studies to modify current practices.

## 5. Conclusions

This study innovatively integrated drug data from different biological, chemical and clinical sources into a prediction model in women’s reproductive health. The model discovers a 2.5% of new DDis potential interactions in the context of women’s reproductive health. Our findings particularly distinguished DDIs that could compromise or boost the efficacy of PCOS treatments, along with novel interactions that may affect contraceptive use, HIV or COVID-19 treatments. Despite the need to experimentally validate the predicted DDIs, these findings elucidate the complexities of drug interactions and provide opportunities for clinically relevant studies.

## Author contributions

P.G.-A.: software, investigation, formal analysis, visualization, writing – original draft, writing – review and edition; I.H.-C.: software, investigation, formal analysis, visualization, writing – original draft, writing – review and edition; P.S.-L.: software, visualization, supervision, writing – review and edition; A.P.-L.: software, data curation; JA.G.-V.: conceptualization, – original draft, writing – review and edition. P.D.-G.: conceptualization, methodology, supervision, project administration, funding acquisition – original draft, writing – review and edition.

---

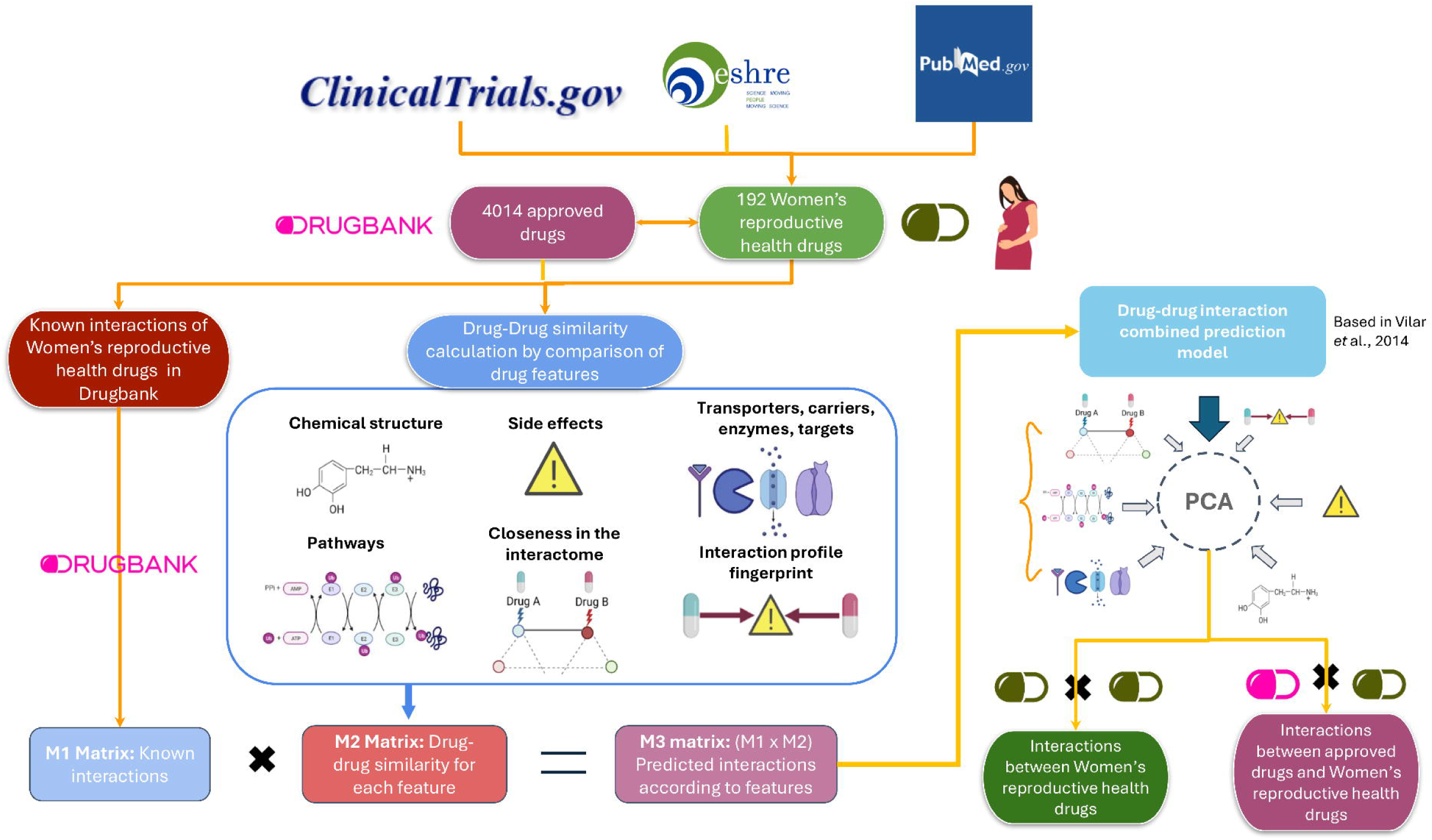

## References

1. Hua ASP, Kincaid Smith P. A comparison of the effects of chlorothiazide and of metolazone in the treatment of hypertension. Clinical Science and Molecular Medicine 1976;51(sup.3):627–9.

2. Becker ML, Kallewaard M, Caspers PWJ, Visser LE, Leufkens HGM, Stricker BHC. Hospitalisations and emergency department visits due to drug-drug interactions: A literature review. Pharmacoepidemiology and Drug Safety 2007;16(6):641–51.

3. Zakharov AV, Varlamova EV, Lagunin AA, Dmitriev AV, Muratov EN, Fourches D, et al. QSAR Modeling and Prediction of Drug-Drug Interactions. Molecular Pharmaceutics 2016;13(2):545–56.

4. Hales CM, Servais J, Martin CB, Kohen D. Prescription Drug Use Among Adults Aged 40-79 in the United States and Canada. NCHS data brief 2019;(347):1–8.

5. Guthrie B, Makubate B, Hernandez-Santiago V, Dreischulte T. The rising tide of polypharmacy and drug-drug interactions: Population database analysis 1995-2010. BMC Medicine 2015;13(1):74.

6. Scarsi KK, Darin KM, Chappell CA, Nitz SM, Lamorde M. Drug–Drug Interactions, Effectiveness, and Safety of Hormonal Contraceptives in Women Living with HIV. Drug Safety 2016;39(11):1053–72.

7. Nanda K, Stuart GS, Robinson J, Gray AL, Tepper NK, Gaffield ME. Drug interactions between hormonal contraceptives and antiretrovirals. Aids 2017;31(7):917–52.

8. Sabers A. Pharmacokinetic interactions between contraceptives and antiepileptic drugs. Seizure 2008;17(2):141–4.

9. Sun H, Sivasubramanian R, Vaidya S, Barve A, Jarugula V. Drug-Drug Interaction Studies With Oral Contraceptives: Pharmacokinetic/Pharmacodynamic and Study Design Considerations. Journal of Clinical Pharmacology 2020;60(S2):S49–62.

10. Lesko LJ, Vozmediano V, Brown JD, Winterstein A, Zhao P, Lippert J, et al. Establishing a Multidisciplinary Framework to Study Drug-Drug Interactions of Hormonal Contraceptives: An Invitation to Collaborate. CPT: Pharmacometrics and Systems Pharmacology 2018;7(11):706–8.

11. Sidra S, Tariq MH, Farrukh MJ, Mohsin M. Evaluation of clinical manifestations, health risks, and quality of life among women with polycystic ovary syndrome. PLoS ONE 2019;14(10):e0223329.

12. Gemmell LC, Williams Z, Forman EJ. Considerations on the restriction of Assisted Reproductive Technology (ART) due to COVID-19. Seminars in Perinatology [Internet] 2020 [cited 2021 Mar 26];44(7). Available from: https://pubmed.ncbi.nlm.nih.gov/33317710/

13. Batiha O, Al-Deeb T, Al-zoubi E, Alsharu E. Impact of COVID-19 and other viruses on reproductive health. Andrologia [Internet] 2020 [cited 2021 Mar 26];52(9). Available from: https://pubmed.ncbi.nlm.nih.gov/32790205/

14. Gerber W, Steyn JD, Kotzé AF, Hamman JH. Beneficial pharmacokinetic drug interactions: A tool to improve the bioavailability of poorly permeable drugs. Pharmaceutics 2018;10(3):106.

15. Vilar S, Harpaz R, Uriarte E, Santana L, Rabadan R, Friedman C. Drug-drug interaction through molecular structure similarity analysis. Journal of the American Medical Informatics Association 2012;19(6):1066–74.

16. Zhang W, Chen Y, Liu F, Luo F, Tian G, Li X. Predicting potential drug-drug interactions by integrating chemical, biological, phenotypic and network data. BMC Bioinformatics 2017;18(1):18.

17. Tatonetti NP, Ye PP, Daneshjou R, Altman RB. Data-driven prediction of drug effects and interactions. Science Translational Medicine 2012;4(125):125ra31.

18. Vilar S, Uriarte E, Santana L, Lorberbaum T, Hripcsak G, Friedman C, et al. Similarity-based modeling in large-scale prediction of drug-drug interactions. Nature Protocols 2014;9(9):2147–63.

19. Ferdousi R, Safdari R, Omidi Y. Computational prediction of drug-drug interactions based on drugs functional similarities. Journal of Biomedical Informatics 2017;70:54–64.

20. Takarabe M, Shigemizu D, Kotera M, Goto S, Kanehisa M. Network-based analysis and characterization of adverse drug-drug interactions. Journal of Chemical Information and Modeling 2011;51(11):2977–85.

21. Park K, Kim D, Ha S, Lee D. Predicting pharmacodynamic drug-drug interactions through signaling propagation interference on protein-protein interaction networks. PLoS ONE 2015;10(10):e0140816.

22. Dunselman GAJ, Vermeulen N, Becker C, Calhaz-Jorge C, D’Hooghe T, De Bie B, et al. ESHRE guideline: Management of women with endometriosis. Human Reproduction 2014;29(3):400–12.

23. Webber L, Davies M, Anderson R, Bartlett J, Braat D, Cartwright B, et al. ESHRE Guideline: Management of women with premature ovarian insufficiency. Human Reproduction 2016;31(5):926–37.

24. Bender Atik R, Christiansen OB, Elson J, Kolte AM, Lewis S, Middeldorp S, et al. ESHRE guideline: recurrent pregnancy loss. Human Reproduction Open 2018;2018(2):1–12.

25. Teede HJ, Misso ML, Costello MF, Dokras A, Laven J, Moran L, et al. Recommendations from the international evidence-based guideline for the assessment and management of polycystic ovary syndrome. Human Reproduction 2018;33(9):1602–18.

26. Gravholt CH, Andersen NH, Conway GS, Dekkers OM, Geffner ME, Klein KO, et al. Clinical practice guidelines for the care of girls and women with Turner syndrome: Proceedings from the 2016 Cincinnati International Turner Syndrome Meeting. European Journal of Endocrinology 2017;177(3):G1–70.

27. Bosch E, Broer S, Griesinger G, Grynberg M, Humaidan P, Kolibianakis E, et al. ESHRE guideline: ovarian stimulation for IVF/ICSI†. Human Reproduction Open 2020;2020(2):1–13.

28. D’Angelo A, Panayotidis C, Amso N, Marci R, Matorras R, Onofriescu M, et al. Recommendations for good practice in ultrasound: oocyte pick up†. Human Reproduction Open 2019;2019(4):1–25.

29. Wishart DS, Knox C, Guo AC, Cheng D, Shrivastava S, Tzur D, et al. DrugBank: A knowledgebase for drugs, drug actions and drug targets. Nucleic Acids Research 2008;36(SUPPL. 1):D901–6.

30. Kuhn M, Letunic I, Jensen LJ, Bork P. The SIDER database of drugs and side effects. Nucleic Acids Research 2016;44(D1):D1075–9.

31. Kanehisa M, Goto S. KEGG: Kyoto Encyclopedia of Genes and Genomes. Nucleic Acids Research 2000;28(1):27–30.

32. Huang J, Niu C, Green CD, Yang L, Mei H, Han JDJ. Systematic Prediction of Pharmacodynamic Drug-Drug Interactions through Protein-Protein-Interaction Network. PLoS Computational Biology 2013;9(3):e1002998.

33. Luck K, Kim DK, Lambourne L, Spirohn K, Begg BE, Bian W, et al. A reference map of the human binary protein interactome. Nature 2020;580(7803):402–8.

34. Bajusz D, Rácz A, Héberger K. Why is Tanimoto index an appropriate choice for fingerprint-based similarity calculations? Journal of Cheminformatics 2015;7(1):20.

35. Core R Team. A Language and Environment for Statistical Computing [Internet]. R Foundation for Statistical Computing. 2019;2:https://www.R--project.org. Available from: http://www.r-project.org

36. Leone Roberti Maggiore U, Scala C, Remorgida V, Venturini PL, Del Deo F, Torella M, et al. Triptorelin for the treatment of endometriosis. Expert Opin Pharmacother 2014;15(8):1153–79.

37. Zhu L, Guan Z, Huang Y, Hua K, Ma L, Zhang J, et al. The efficacy and safety of triptorelin-therapy following conservative surgery for deep infiltrating endometriosis: A multicenter, prospective, non-interventional study in China. Medicine 2022;101(5):e28766.

38. Acmaz G, Cinar L, Acmaz B, Aksoy H, Kafadar YT, Madendag Y, et al. The Effects of Oral Isotretinoin in Women with Acne and Polycystic Ovary Syndrome. BioMed Research International [Internet] 2019 [cited 2021 Sep 8];2019. Available from: https://pubmed.ncbi.nlm.nih.gov/31080813/

39. Dungan HM, Clifton DK, Steiner RA. Minireview: Kisspeptin neurons as central processors in the regulation of gonadotropin-releasing hormone secretion. Endocrinology 2006;147(3):1154–8.

40. Di Emidio G, Rea F, Placidi M, Rossi G, Cocciolone D, Virmani A, et al. Regulatory functions of l-carnitine, acetyl, and propionyl l-carnitine in a PCOS mouse model: Focus on antioxidant/antiglycative molecular pathways in the ovarian microenvironment. Antioxidants 2020;9(9):1–17.

41. El Sharkwy I, Sharaf El-Din M. l-Carnitine plus metformin in clomiphene-resistant obese PCOS women, reproductive and metabolic effects: a randomized clinical trial. Gynecological Endocrinology 2019;35(8):701–5.

42. Knight JA, Wong J, Blackmore KM, Raboud JM, Vieth R. Vitamin D association with estradiol and progesterone in young women. Cancer Causes and Control 2010;21(3):479–83.

43. Freedman DM, Looker AC, Chang SC, Graubard BI. Prospective study of serum vitamin D and cancer mortality in the United States. Journal of the National Cancer Institute 2007;99(21):1594–602.

44. Chantalat E, Valera M-C, Vaysse C, Noirrit E, Rusidze M, Weyl A, et al. Estrogen Receptors and Endometriosis. International Journal of Molecular Sciences 2020;21(8):2815.

45. Lu Y, Chen R, Cai J, Huang Z, Yuan H. The management of hypertension in women planning for pregnancy. British Medical Bulletin 2018;128(1):75–84.

46. Lai SF, Lam MT, Li HWR, Nga EHY. A randomized double-blinded non-inferiority trial comparing fentanyl and midazolam with pethidine and diazepam for pain relief during oocyte retrieval. Reproductive BioMedicine Online 2020;40(5):653–60.

47. Li L, Zhang R, Zeng J, Ke H, Peng X, Huang L, et al. Effectiveness and safety assessment of drospirenone/ethinyl estradiol tablet in treatment of PCOS patients: A single center, prospective, observational study. BMC Women’s Health [Internet] 2020 [cited 2021 Jun 29];20(1). Available from: https://pubmed.ncbi.nlm.nih.gov/32106860/

48. Lin YH, Huang MZ, Hwang JL, Chen HJ, Hsieh BC, Huang LW, et al. Combination of cabergoline and embryo cryopreservation after GnRH agonist triggering prevents OHSS in patients with extremely high estradiol levels - A retrospective study. Journal of Assisted Reproduction and Genetics 2013;30(6):753–9.

49. Zhang W, Tian Y, Xie D, Miao Y, Liu J, Wang X. The impact of peak estradiol during controlled ovarian stimulation on the cumulative live birth rate of IVF/ICSI in non-PCOS patients. Journal of Assisted Reproduction and Genetics 2019;36(11):2333–44.

50. Ullah K, Rahman TU, Pan HT, Guo MX, Dong XY, Liu J, et al. Serum estradiol levels in controlled ovarian stimulation directly affect the endometrium. Journal of Molecular Endocrinology 2017;59(2):105–19.

51. Pinheiro LMA, Cândido P da S, Moreto TC, Di Almeida WG, De Castro EC. Estradiol use in the luteal phase and its effects on pregnancy rates in IVF cycles with GnRH antagonist: A systematic review. Jornal Brasileiro de Reproducao Assistida 2017;21(3):247–50.

52. Labarta E, Rodríguez C. Progesterone use in assisted reproductive technology. Best Practice and Research: Clinical Obstetrics and Gynaecology 2020;69:74–84.

53. Eichner E. Progestins. The American journal of nursing 1965;65:78–81.

54. Plewig G, Kligman AM. Estrogens and Oral Contraceptives. ACNE and ROSACEA 1993;659–61.

55. Sundaram V, Björnsson ES. Drug-induced cholestasis. Hepatology Communications 2017;1(8):726–35.

56. Tivoli YA, Rubenstein RM. Pruritus: An updated look at an old problem. Journal of Clinical and Aesthetic Dermatology 2009;2(7):30–6.

57. Xie C, Aloreidi K, Quist E, Pocha C. A Rare Cause of Hepatomegaly: Oral Contraceptives-Induced Peliosis Hepatis: 2255. Official journal of the American College of Gastroenterology | ACG 2017;112:S1240.

58. Wang S, Wang Y, Xu J, Chen Y. Is the oral contraceptive or hormone replacement therapy a risk factor for cholelithiasis. Medicine (United States) 2017;96(14):e6556.

59. Batta A, Kalra B, Khirasaria R. Trends in FDA drug approvals over last 2 decades: An observational study. Journal of Family Medicine and Primary Care 2020;9(1):105.

