## Supplemental Table 1 for "Identification of novel drug-drug interactions: towards precision medicine in women’s reproductive health"

**Supplemental Table 1. Classification of reproductive categories and drugs currently used in women’s reproductive health. (A)** Keywords used to search the ESHRE guidelines, DrugBank, and ClinicalTrials.gov were classified into seven reproductive categories. **(B)** A total of 192 drugs currently prescribed for female reproduction indications were retrieved from ClinicalTrials.gov, DrugBank, ESHRE guidelines, and PubMed. Drugs were classified into one or more of the corresponding reproductive categories presented in (A), according to the associated keywords in the ESHRE guidelines or ClinicalTrials.gov. COS, controlled ovarian stimulation; POI, premature ovarian insufficiency; POF, premature ovarian failure; PCOS, polycystic ovary syndrome; RPL, recurrent pregnancy loss.

**(A)**

| **Reproductive category** | **ESHRE**  **Guideline(s)** | **DrugBank keywords** | **ClinicalTrials*.*gov keywords** |
| --- | --- | --- | --- |
| **Preconception** | - | Contraceptives | Contraception; Contraceptive usage; Contraceptive, complications, intrauterine; Contraceptive, complications, intrauterine, mechanical |
| **Infertility & IVF** | COS; Oocyte retrieval | Infertility | Implantation; Endometrial receptivity; Reproductive Techniques; Reproductive Sterility; Infertility, Female; Infertility Unexplained; Infertility of Tubal Origin; Infertility Secondary; Ovarian Stimulation; Controlled Ovarian Stimulation; Fertility disorders; Fertility Issues; Subfertility; Assisted Reproductive Technology; Assisted Reproductive; Assisted Reproduction |
| **Menopause** | - | - | Hormone Replacement Therapy |
| **Uterine disease** | Endometriosis | - | Endometriosis; Endometrial diseases; Uterus Prolapse; Uterine Fibroid; Uterus Disease; Uterine Bleeding; Uterus Abnormal; Asherman’s Syndrome |
| **Ovarian disease** | POI; PCOS | - | Ovulation Disorder; Ovulation Absent; Ovulation, Failure or Lack of; Anovulation; POF; PCOS |
| **Menstrual disorder** | - | - | Amenorrhea; Amenorrhea Traumatica; Dysmenorrhea |
| **Other reproductive disorder** | RPL; Turner Syndrome | - | Pregnancy Loss; Recurrent Pregnancy Loss; Turner Syndrome; Gynecologic Disease; Reproductive Disorder; Reproductive System Disorder; Preeclampsia; X Fragile Syndrome; Hyperprolactinemia |

**(B)**

| ID | Drug name | Reproductive category | ESHRE guideline(s) | ClinicalTrials.gov keywords |
| --- | --- | --- | --- | --- |
| DB00284 | Acarbose | Ovarian diseases | - | PCOS |
| DB00316 | Acetaminophen | Uterus diseases - Infertility & IVF - Menstrual disorders - Other reproductive diseases | Endometriosis; Oocyte retrieval | Uterine Fibroid; Dysmenorrhea; Preeclampsia; Uterus Disease; Fertility Issues; Infertility, Female; Pregnancy loss |
| DB00945 | Acetylsalicylic acid | Other reproductive diseases | RPL | Preeclampsia; Recurrent Pregnancy Loss; Pregnancy loss |
| DB06203 | Alogliptin | Ovarian diseases | - | PCOS |
| DB00321 | Amitriptyline | Uterus diseases | Endometriosis | - |
| DB01060 | Amoxicillin | Uterus diseases | - | Uterus Disease |
| DB01217 | Anastrozole | Uterus diseases - Ovarian diseases | Endometriosis; PCOS | Uterine Fibroid; Uterine bleeding; Uterus Disease; Endometriosis |
| DB01238 | Aripiprazole | Other reproductive diseases | - | Hyperprolactinemia |
| DB01076 | Atorvastatin | Ovarian diseases | PCOS | - |
| DB00207 | Azithromycin | Other reproductive diseases - Uterus diseases | - | Preeclampsia; Uterus Disease |
| DB06401 | Bazedoxifene | Menopause - Other reproductive diseases | - | Hormone Replacement Therapy; Reproductive System Disorder |
| DB09258 | Bemiparin | Infertility & IVF | - | Assisted Reproductive Technology; Assisted Reproductive; Fertility Issues; Reproductive Sterility |
| DB11105 | Benzalkonium | Preconception | - | Contraception |
| DB01200 | Bromocriptine | Other reproductive diseases - Infertility & IVF | RPL; Infertility | - |
| DB00297 | Bupivacaine | Uterus diseases | - | Uterine Fibroid; Uterus Prolapse; Uterus Disease |
| DB00921 | Buprenorphine | Other reproductive diseases | - | Pregnancy loss |
| DB06719 | Buserelin | Uterus diseases - Ovarian diseases - Infertility & IVF | Endometriosis; PCOS; COS | Uterus Disease; Fertility Issues; Ovarian Stimulation |
| DB00248 | Cabergoline | Infertility & IVF | - | Fertility Issues; Infertility, Female |
| DB06724 | Calcium carbonate | Menstrual disorders - Uterus diseases | - | Amenorrhea; Endometriosis |
| DB01197 | Captopril | Other reproductive diseases | - | Preeclampsia |
| DB01282 | Carbetocin | Uterus diseases | - | Uterine Fibroid; Uterine bleeding |
| DB01327 | Cefazolin | Uterus diseases | - | Uterine bleeding; Uterus Disease |
| DB00050 | Cetrorelix | Ovarian diseases - Infertility & IVF | PCOS; COS | Assisted Reproduction; Assisted Reproductive; Fertility Issues; Infertility, Female; Controlled Ovarian Stimulation; Endometrial receptivity; Ovarian Stimulation; PCOS; Reproductive Sterility |
| DB01161 | Chloroprocaine | Uterus diseases | - | Uterus Disease |
| DB00169 | Cholecalciferol | Menstrual disorders - Other reproductive diseases - Uterus diseases | - | Amenorrhea; Preeclampsia; Endometriosis |
| DB00097 | Choriogonadotropin alfa | Infertility & IVF - Ovarian diseases | COS; Infertility | Fertility Issues; Infertility Unexplained; Ovarian Stimulation; PCOS |
| DB09126 | Chorionic Gonadotropin (Human) | Uterus diseases - Infertility & IVF - Menopause - Ovarian diseases - Other reproductive diseases | Endometriosis; COS | Assisted Reproduction; Assisted Reproductive Technology; Assisted Reproductive; Fertility Issues; Hormone Replacement Therapy; Infertility, Female; Subfertility; Controlled Ovarian Stimulation; Ovarian Stimulation; PCOS; Pregnancy loss |
| DB04272 | Citric acid | Preconception | Contraceptives | - |
| DB01190 | Clindamycin | Uterus diseases - Other reproductive diseases | - | Uterine bleeding; Uterus Disease; Pregnancy loss |
| DB00882 | Clomifene | Uterus diseases - Ovarian diseases - Infertility & IVF - Menopause | Endometriosis; PCOS; COS; Infertility | Assisted Reproductive; Fertility Issues; Hormone Replacement Therapy; Infertility, Female; Subfertility; Controlled Ovarian Stimulation; Endometriosis; Infertility Unexplained; Ovarian Stimulation; Ovulation Disorder; Ovulation, Absent; PCOS; Reproductive Sterility |
| DB00575 | Clonidine | Other reproductive diseases | - | Preeclampsia |
| DB00758 | Clopidogrel | Infertility & IVF | - | Fertility Issues; Infertility, Female |
| DB00318 | Codeine | Infertility & IVF | Oocyte retrieval | - |
| DB00286 | Conjugated estrogens | Ovarian diseases - Other reproductive diseases - Uterus diseases - Menopause | POI; Turner syndrome | Uterus Prolapse; Uterus Disease; Hormone Replacement Therapy; Reproductive System Disorder; Endometrial diseases |
| DB09130 | Copper | Preconception | Contraceptives | Contraception |
| DB09066 | Corifollitropin alfa | Infertility & IVF - Uterus diseases - Ovarian diseases | COS | Uterus Disease; Fertility Issues; Infertility, Female; Controlled Ovarian Stimulation; Endometrial diseases; Endometrial receptivity; Ovarian Stimulation; PCOS; Reproductive Sterility |
| DB11672 | Curcumin | Preconception - Uterus diseases | - | Contraceptive Usage; Uterine bleeding; Uterus Disease |
| DB04839 | Cyproterone acetate | Uterus diseases - Ovarian diseases - Preconception - Other reproductive diseases | Endometriosis; PCOS; Contraceptives | Reproductive System Disorder; PCOS |
| DB01406 | Danazol | Uterus diseases | Endometriosis | - |
| DB00304 | Desogestrel | Uterus diseases - Ovarian diseases - Preconception - Other reproductive diseases - Menstrual disorders - Infertility & IVF | Endometriosis; PCOS; Contraceptives; Turner syndrome | Dysmenorrhea; Fertility Issues; Ovarian Stimulation |
| DB01234 | Dexamethasone | Ovarian diseases - Infertility & IVF | PCOS | Fertility Issues |
| DB09214 | Dexketoprofen | Menstrual disorders | - | Dysmenorrhea |
| DB00829 | Diazepam | Infertility & IVF | - | Fertility Issues |
| DB00586 | Diclofenac | Infertility & IVF - Menstrual disorders | Oocyte retrieval | Dysmenorrhea; Fertility Issues; Infertility, Female; Infertility of Tubal Origin |
| DB09123 | Dienogest | Uterus diseases - Ovarian diseases - Preconception - Menstrual disorders - Infertility & IVF | Endometriosis; PCOS ;Contraceptives | Uterine Fibroid; Dysmenorrhea; Contraceptive Usage; Contraception; Uterus Disease; Fertility Issues; Endometriosis |
| DB00255 | Diethylstilbestrol | Preconception | Contraceptives | - |
| DB00390 | Digoxin | Other reproductive diseases | - | Pregnancy loss |
| DB00917 | Dinoprostone | Preconception - Infertility & IVF | - | Contraception; Fertility Issues |
| DB00470 | Dronabinol | Other reproductive diseases | - | Pregnancy loss |
| DB01395 | Drospirenone | Ovarian diseases - Preconception - Menstrual disorders - Uterus diseases | PCOS; Contraceptives | Dysmenorrhea; Contraception; Endometriosis; PCOS |
| DB00476 | Duloxetine | Uterus diseases | Endometriosis | - |
| DB00378 | Dydrogesterone | Uterus diseases - Ovarian diseases - Preconception - Infertility & IVF - Menopause - Other reproductive diseases | Endometriosis; POI; PCOS; Contraceptives; COS; Infertility | Uterus Disease; Fertility Issues; Hormone Replacement Therapy; Infertility, Female; Reproductive System Disorder; Recurrent Pregnancy Loss; Endometrial diseases; Implantation, placenta; Pregnancy loss; Reproductive Techniques |
| DB06243 | Eflornithine | Ovarian diseases | PCOS | - |
| DB11979 | Elagolix | Uterus diseases - Ovarian diseases | Endometriosis | Uterine Fibroid; Uterine bleeding; Uterus Disease; Anovulation; Ovulation, Absent; Ovulation, Failure or lack of |
| DB00584 | Enalapril | Other reproductive diseases | - | Preeclampsia |
| DB01225 | Enoxaparin | Infertility & IVF - Other reproductive diseases | - | Assisted Reproductive Technology; Assisted Reproductive; Fertility Issues; Infertility, Female; Infertility Unexplained; Pregnancy loss |
| DB01253 | Ergometrine | Uterus diseases | - | Uterine bleeding |
| DB01175 | Escitalopram | Ovarian diseases | - | PCOS |
| DB09381 | Esterified estrogens | Ovarian diseases | POI | - |
| DB00783 | Estradiol | Uterus diseases - Ovarian diseases - Preconception - Other reproductive diseases - Infertility & IVF - Menopause | Endometriosis; POI; PCOS; Contraceptives; Turner syndrome | Contraception; Uterine bleeding; Uterus Disease; Fertility Issues; Hormone Replacement Therapy; Reproductive System Disorder; Pregnancy loss |
| DB13952 | Estradiol acetate | Ovarian diseases | - | PCOS |
| DB13954 | Estradiol cypionate | Preconception | Contraceptives | - |
| DB13956 | Estradiol valerate | Ovarian diseases - Preconception - Menstrual disorders - Uterus diseases - Infertility & IVF - Menopause - Other reproductive diseases | POI; Contraceptives | Amenorrhea Traumatica; Dysmenorrhea; Asherman’s Syndrome; Contraception; Fertility Issues; Hormone Replacement Therapy; Infertility, Female; Endometrial diseases; Endometriosis; Ovarian Stimulation; Ovulation Disorder; Ovulation, Absent; PCOS; Pregnancy loss |
| DB04573 | Estriol | Uterus diseases - Other reproductive diseases | - | Uterus Prolapse; Uterus Disease; Reproductive System Disorder; Endometrial diseases |
| DB00977 | Ethinylestradiol | Uterus diseases - Ovarian diseases - Preconception - Other reproductive diseases - Menstrual disorders | Endometriosis; PCOS; Contraceptives; Turner syndrome | Uterus Abnormal; Amenorrhea; Dysmenorrhea; Contraceptive Usage; Contraception; Uterine bleeding; Uterus Disease; Reproductive System Disorder ;Endometriosis; PCOS |
| DB00965 | Ethiodized oil | Infertility & IVF | - | Fertility Issues; Infertility, Female |
| DB00823 | Ethynodiol diacetate | Uterus diseases - Ovarian diseases - Preconception | Endometriosis; PCOS; Contraceptives | - |
| DB00292 | Etomidate | Other reproductive diseases | - | Pregnancy loss |
| DB00294 | Etonogestrel | Uterus diseases - Preconception - Ovarian diseases - Other reproductive diseases | Endometriosis; Contraceptives | Contraceptive Usage; Contraception; Uterine bleeding; Uterus Disease; Endometriosis; PCOS; Pregnancy loss |
| DB00990 | Exemestane | Uterus diseases | Endometriosis | - |
| DB01276 | Exenatide | Other reproductive diseases - Ovarian diseases | - | Reproductive System Disorder; PCOS; Reproductive Disorder |
| DB00813 | Fentanyl | Uterus diseases - Infertility & IVF | Endometriosis; Oocyte retrieval | Uterus Disease; Fertility Issues |
| DB08917 | Ferric carboxymaltose | Infertility & IVF | - | Assisted Reproductive Technology; Assisted Reproductive; Fertility Issues |
| DB13257 | Ferrous sulfate anhydrous | Uterus diseases | - | Uterine Fibroid; Uterus Abnormal; Uterine bleeding; Uterus Disease |
| DB06215 | Ferumoxytol | Uterus diseases | - | Uterus Abnormal; Uterine bleeding ;Uterus Disease |
| DB09222 | Fibrinogen human | Uterus diseases | - | Uterine bleeding |
| DB00099 | Filgrastim | Infertility & IVF - Ovarian diseases | - | Fertility Issues; POF |
| DB01216 | Finasteride | Ovarian diseases | PCOS | - |
| DB13146 | Fluciclovine (18F) | Uterus diseases | - | Endometrial diseases |
| DB00499 | Flutamide | Ovarian diseases | PCOS | - |
| DB00158 | Folic acid | Infertility & IVF - Ovarian diseases | - | Fertility Issues; Infertility, Female; PCOS |
| DB00066 | Follitropin | Uterus diseases - Infertility & IVF - Ovarian diseases | Endometriosis; COS; Infertility | Fertility disorders; Assisted Reproduction; Assisted Reproductive Technology; Assisted Reproductive; Uterus Disease; Fertility Issues; Infertility, Female; Subfertility; Anovulation; Controlled Ovarian Stimulation; Endometrial diseases; Endometrial receptivity; Ovarian Stimulation; Ovulation, Absent; PCOS; Reproductive Sterility; Reproductive Techniques |
| DB00695 | Furosemide | Other reproductive diseases | - | Preeclampsia |
| DB00996 | Gabapentin | Uterus diseases - Other reproductive diseases | Endometriosis | Pregnancy loss |
| DB06785 | Ganirelix | Infertility & IVF | COS | Assisted Reproductive Technology ;Assisted Reproductive; Fertility Issues; Controlled Ovarian Stimulation; Ovarian Stimulation; Reproductive Sterility |
| DB11619 | Gestrinone | Uterus diseases - Preconception - Infertility & IVF | Endometriosis; Contraceptives; Infertility | - |
| DB00040 | Glucagon | Infertility & IVF | - | Fertility disorders |
| DB00644 | Gonadorelin | Infertility & IVF - Ovarian diseases | COS; Infertility | Fertility Issues ;Ovulation Disorder; PCOS |
| DB00014 | Goserelin | Uterus diseases | Endometriosis | Uterine Fibroid; Uterine bleeding; Uterus Disease |
| DB01109 | Heparin | Other reproductive diseases | RPL | Recurrent Pregnancy Loss; Pregnancy loss |
| DB00028 | Human immunoglobulin G | Other reproductive diseases | - | Pregnancy loss |
| DB06789 | Hydroxyprogesterone caproate | Preconception - Other reproductive diseases | Contraceptives | Preeclampsia |
| DB01050 | Ibuprofen | Uterus diseases - Other reproductive diseases | Endometriosis | Uterine Fibroid; Preeclampsia; Uterus Disease; Pregnancy loss |
| DB09374 | Indocyanine green acid form | Uterus diseases | - | Endometriosis |
| DB13178 | Inositol | Ovarian diseases - Infertility & IVF | PCOS; Infertility | Assisted Reproduction; Assisted Reproductive Technology; Assisted Reproductive; Infertility, Female; PCOS; Reproductive Sterility |
| DB00951 | Isoniazid | Infertility & IVF - Other reproductive diseases | - | Contraceptive; Complications, Intrauterine, Mechanical; Fertility Issues; Infertility, Female; Pregnancy loss |
| DB01020 | Isosorbide mononitrate | Other reproductive diseases | - | Pregnancy loss |
| DB00982 | Isotretinoin | Ovarian diseases | - | PCOS |
| DB01026 | Ketoconazole | Ovarian diseases | PCOS | - |
| DB00465 | Ketorolac | Preconception - Uterus diseases | - | Contraception; Uterine bleeding |
| DB00598 | Labetalol | Other reproductive diseases | - | Preeclampsia |
| DB04398 | Lactic acid | Preconception | Contraceptives | - |
| DB01006 | Letrozole | Uterus diseases - Ovarian diseases - Infertility & IVF - Other reproductive diseases | Endometriosis; PCOS; COS | Uterine Fibroid; Fertility Issues; Infertility, Female; Subfertility; Anovulation ;Controlled Ovarian Stimulation ;Endometrial diseases; Endometrial receptivity; Ovarian Stimulation; Ovulation Disorder; Ovulation, Absent; PCOS; Pregnancy loss; Reproductive Techniques |
| DB00007 | Leuprolide | Uterus diseases - Ovarian diseases - Infertility & IVF - Menstrual disorders - Preconception - Other reproductive diseases | Endometriosis; PCOS; COS | Uterine Fibroid; Amenorrhea; Assisted Reproductive Technology; Assisted Reproductive; Contraception; Fertility Issues; Endometriosis; Ovarian Stimulation; PCOS; Pregnancy loss |
| DB01002 | Levobupivacaine | Other reproductive diseases | - | Preeclampsia |
| DB00583 | Levocarnitine | Ovarian diseases | - | PCOS |
| DB00367 | Levonorgestrel | Uterus diseases - Ovarian diseases - Preconception - Other reproductive diseases - Menstrual disorders | Endometriosis; PCOS; Contraceptives; Turner syndrome | Uterine Fibroid; Uterus Abnormal; Amenorrhea; Contraceptive Usage; Contraception; Uterine bleeding; Uterus Disease; Endometriosis |
| DB00281 | Lidocaine | Infertility & IVF - Uterus diseases - Other reproductive diseases - Preconception | Oocyte retrieval | Uterine Fibroid; Uterus Prolapse; Preeclampsia; Contraception; Uterus Disease; Fertility Issues; Infertility, Female; Infertility of Tubal Origin; Pregnancy loss |
| DB00166 | Lipoic acid | Infertility & IVF | - | Fertility Issues; Infertility, Female; Endometrial receptivity |
| DB06655 | Liraglutide | Ovarian diseases - Infertility & IVF - Other reproductive diseases | PCOS | Fertility Issues; Infertility, Female; Reproductive System Disorder; PCOS |
| DB01255 | Lisdexamfetamine | Ovarian diseases | - | POF |
| DB00227 | Lovastatin | Other reproductive diseases | - | X Fragile Syndrome |
| DB14741 | Luteinizing hormone | Infertility & IVF | COS | - |
| DB00044 | Lutropin alfa | Infertility & IVF | COS; Infertility | Assisted Reproductive; Fertility Issues; Infertility, Female; Controlled Ovarian Stimulation; Ovarian Stimulation; Reproductive Sterility |
| DB12474 | Lynestrenol | Preconception | Contraceptives | - |
| DB00653 | Magnesium sulfate | Other reproductive diseases | - | Preeclampsia |
| DB09124 | Medrogestone | Preconception | Contraceptives | - |
| DB00603 | Medroxyprogesterone acetate | Uterus diseases - Ovarian diseases - Preconception - Other reproductive diseases - Infertility & IVF | Endometriosis; POI; PCOS; Contraceptives; Turner syndrome | Contraception; Fertility Issues; Anovulation; Endometrial diseases; Endometriosis; Ovarian Stimulation; Ovulation Disorder; PCOS |
| DB00351 | Megestrol acetate | Uterus diseases - Preconception | Endometriosis; Contraceptives | - |
| DB01065 | Melatonin | Other reproductive diseases | - | Reproductive System Disorder |
| DB00032 | Menotropins | Infertility & IVF | COS; Infertility | Assisted Reproduction; Assisted Reproductive Technology; Assisted Reproductive; Fertility Issues; Infertility, Female; Controlled Ovarian Stimulation; Ovarian Stimulation; Reproductive Sterility |
| DB00454 | Meperidine | Infertility & IVF | - | Fertility Issues |
| DB01357 | Mestranol | Uterus diseases - Ovarian diseases - Preconception | Endometriosis; PCOS; Contraceptives | - |
| DB04817 | Metamizole | Uterus diseases | Endometriosis | - |
| DB00331 | Metformin | Ovarian diseases - Other reproductive diseases - Infertility & IVF - Uterus diseases | PCOS | Preeclampsia; Fertility Issues; Infertility, Female; Reproductive System Disorder; Anovulation; Endometrial diseases; Ovulation, Absent; PCOS; Reproductive Disorder |
| DB00333 | Methadone | Uterus diseases | Endometriosis | - |
| DB00134 | Methionine | Ovarian diseases | - | PCOS |
| DB00968 | Methyldopa | Other reproductive diseases | - | Preeclampsia |
| DB00353 | Methylergometrine | Uterus diseases - Other reproductive diseases | - | Uterine bleeding; Pregnancy loss |
| DB00683 | Midazolam | Infertility & IVF - Other reproductive diseases | Oocyte retrieval | Fertility Issues; Pregnancy loss |
| DB00834 | Mifepristone | Uterus diseases - Preconception - Other reproductive diseases | Endometriosis; Contraceptives | Uterine Fibroid; Contraception; Pregnancy loss |
| DB00929 | Misoprostol | Uterus diseases - Preconception - Other reproductive diseases | - | Uterine Fibroid; Contraceptive Usage; Contraceptive; Complications, Intrauterine; Preeclampsia; Uterine bleeding; Pregnancy loss |
| DB00295 | Morphine | Uterus diseases | Endometriosis | - |
| DB00666 | Nafarelin | Uterus diseases - Infertility & IVF | Endometriosis; COS | - |
| DB00788 | Naproxen | Uterus diseases - Menstrual disorders | Endometriosis | Uterine Fibroid; Dysmenorrhea; Uterine bleeding; Uterus Disease; Endometrial diseases |
| DB01115 | Nifedipine | Other reproductive diseases | - | Preeclampsia |
| DB00727 | Nitroglycerin | Preconception | - | Contraception |
| DB11636 | Nomegestrol | Uterus diseases - Preconception | Endometriosis | Contraception; Uterine bleeding |
| DB06804 | Nonoxynol-9 | Preconception | Contraceptives | - |
| DB06713 | Norelgestromin | Uterus diseases - Ovarian diseases - Preconception | Endometriosis; PCOS; Contraceptives | - |
| DB00717 | Norethisterone | Uterus diseases - Ovarian diseases - Preconception - Other reproductive diseases | Endometriosis; PCOS; Contraceptives; Turner syndrome | Contraception; Uterine bleeding; Uterus Disease; Endometriosis |
| DB09371 | Norethynodrel | Preconception | Contraceptives | - |
| DB00957 | Norgestimate | Ovarian diseases - Preconception - Other reproductive diseases | PCOS; Contraceptives; Turner syndrome | - |
| DB09389 | Norgestrel | Uterus diseases - Ovarian diseases - Preconception | Endometriosis; PCOS; Contraceptives | - |
| DB01083 | Orlistat | Ovarian diseases | - | PCOS |
| DB00621 | Oxandrolone | Other reproductive diseases | Turner syndrome | - |
| DB00497 | Oxycodone | Other reproductive diseases | - | Pregnancy loss |
| DB00107 | Oxytocin | Uterus diseases - Other reproductive diseases | - | Uterine Fibroid; Preeclampsia; Uterine bleeding; Uterus Disease; Pregnancy loss |
| DB01267 | Paliperidone | Other reproductive diseases | - | Hyperprolactinemia |
| DB00652 | Pentazocine | Other reproductive diseases | - | Pregnancy loss |
| DB01132 | Pioglitazone | Ovarian diseases | PCOS | PCOS |
| DB00554 | Piroxicam | Menstrual disorders | - | Dysmenorrhea |
| DB13968 | Plasma protein fraction (human) | Uterus diseases | - | Uterine bleeding |
| DB06812 | Povidone-iodine | Uterus diseases | - | Uterus Disease |
| DB01708 | Prasterone | Preconception - Infertility & IVF | - | Contraceptive Usage; Contraception; Fertility Issues; Subfertility; Reproductive Sterility |
| DB00860 | Prednisolone | Infertility & IVF - Ovarian diseases - Other reproductive diseases | - | Fertility Issues; Infertility, Female; PCOS; Pregnancy loss |
| DB00635 | Prednisone | Other reproductive diseases | - | Recurrent Pregnancy Loss; Pregnancy loss |
| DB00230 | Pregabalin | Uterus diseases - Other reproductive diseases | Endometriosis | Pregnancy loss |
| DB00750 | Prilocaine | Preconception - Uterus diseases | - | Contraception; Uterus Disease |
| DB00396 | Progesterone | Uterus diseases - Ovarian diseases - Preconception - Infertility & IVF - Other reproductive diseases - Menopause | Endometriosis; POI; PCOS; Contraceptives; COS; Turner syndrome; Infertility | Assisted Reproduction; Assisted Reproductive Technology; Assisted Reproductive; Contraception; Uterus Disease; Fertility Issues; Hormone Replacement Therapy; Infertility, Female; Reproductive System Disorder; Recurrent Pregnancy Loss; Endometrial receptivity; Implantation, placenta; Ovarian Stimulation; Pregnancy loss; Reproductive Sterility; Reproductive Techniques |
| DB00818 | Propofol | Ovarian diseases - Other reproductive diseases | - | PCOS; Pregnancy loss |
| DB00165 | Pyridoxine | Uterus diseases | Endometriosis | - |
| DB01045 | Rifampicin | Preconception | - | Contraception |
| DB00734 | Risperidone | Other reproductive diseases | - | Hyperprolactinemia |
| DB00728 | Rocuronium | Uterus diseases - Other reproductive diseases | - | Uterus Prolapse; Gynecologic Disease; Uterus Disease; Endometriosis |
| DB00533 | Rofecoxib | Uterus diseases | Endometriosis | - |
| DB01656 | Roflumilast | Ovarian diseases | - | PCOS |
| DB00412 | Rosiglitazone | Ovarian diseases | PCOS | - |
| DB01098 | Rosuvastatin | Ovarian diseases | PCOS | - |
| DB14583 | Segesterone acetate | Preconception | Contraceptives | - |
| DB13928 | Semaglutide | Ovarian diseases | - | PCOS |
| DB00203 | Sildenafil | Infertility & IVF | - | Fertility Issues |
| DB00641 | Simvastatin | Ovarian diseases | PCOS | PCOS |
| DB00877 | Sirolimus | Uterus diseases | - | Uterine Fibroid |
| DB01261 | Sitagliptin | Ovarian diseases | - | PCOS |
| DB00052 | Somatotropin | Other reproductive diseases - Infertility & IVF | Turner syndrome | Assisted Reproduction; Assisted Reproductive Technology; Assisted Reproductive; Fertility Issues; Infertility, Female; Subfertility; Turner Syndrome |
| DB09422 | Soybean oil | Other reproductive diseases | - | Recurrent Pregnancy Loss; Pregnancy loss |
| DB00421 | Spironolactone | Ovarian diseases - Preconception | PCOS; Contraceptives | - |
| DB06206 | Sugammadex | Other reproductive diseases | - | Gynecologic Disease |
| DB00675 | Tamoxifen | Infertility & IVF - Preconception - Uterus diseases - Ovarian diseases | COS | Contraception; Uterine bleeding; Uterus Disease; Fertility Issues; PCOS |
| DB00624 | Testosterone | Infertility & IVF | - | Fertility Issues; Infertility, Female |
| DB00152 | Thiamine | Uterus diseases | Endometriosis | - |
| DB09070 | Tibolone | Uterus diseases - Menstrual disorders | Endometriosis | Amenorrhea; Endometriosis |
| DB00273 | Topiramate | Ovarian diseases | - | PCOS |
| DB00193 | Tramadol | Menstrual disorders | - | Dysmenorrhea |
| DB00302 | Tranexamic acid | Uterus diseases | - | Uterine Fibroid; Uterine bleeding; Uterus Disease |
| DB06825 | Triptorelin | Uterus diseases - Ovarian diseases - Infertility & IVF | Endometriosis; PCOS; COS | Assisted Reproduction; Assisted Reproductive Technology; Assisted Reproductive; Uterus Disease; Fertility Issues; Infertility, Female; Controlled Ovarian Stimulation; Endometriosis; Ovarian Stimulation ;PCOS; Reproductive Sterility |
| DB08867 | Ulipristal | Preconception - Uterus diseases - Menstrual disorders - Infertility & IVF - Other reproductive diseases | Contraceptives | Uterine Fibroid; Dysmenorrhea; Contraception; Uterine bleeding; Uterus Disease; Fertility Issues; Infertility, Female; Endometriosis; Pregnancy loss |
| DB00094 | Urofollitropin | Infertility & IVF | COS; Infertility | Assisted Reproduction; Assisted Reproductive; Fertility Issues |
| DB00067 | Vasopressin | Uterus diseases | - | Uterine Fibroid |
| DB11094 | Vitamin D | Other reproductive diseases - Infertility & IVF - Menopause - Ovarian diseases | Turner syndrome | Fertility Issues; Hormone Replacement Therapy; Infertility, Female; PCOS |
| DB00682 | Warfarin | Other reproductive diseases | - | Pregnancy loss |
