## Supplemental Table 4 for "Identification of novel drug-drug interactions: towards precision medicine in women’s reproductive health"

| Disease | | DDI | WRHD | Predicted Effect |
| --- | --- | --- | --- | --- |
| HIV | Amphotericin B-Ethinylestradiol | | Ethinylestradiol | Amphotericin B may decrease the excretion rate of Ethinylestradiol which could result in a higher serum level. |
| HIV | Amphotericin B-Mifepristone | | Mifepristone | The metabolism of Mifepristone can be decreased when combined with Amphotericin B. |
| HIV | Amphotericin B-Mestranol | | Mestranol | Amphotericin B may decrease the excretion rate of Mestranol which could result in a higher serum level. |
| HIV | Desogestrel-Amphotericin B | | Desogestrel | The metabolism of Amphotericin B can be increased when combined with Desogestrel. |
| HIV | Amphotericin B-Segesterone acetate | | Segesterone acetate | The metabolism of Segesterone acetate can be decreased when combined with Amphotericin B. |
| HIV | Amphotericin B-Norethynodrel | | Norethynodrel | The metabolism of Norethynodrel can be decreased when combined with Amphotericin B. |
| HIV | Levonorgestrel-Amphotericin B | | Levonorgestrel | The metabolism of Amphotericin B can be decreased when combined with Levonorgestrel. |
| HIV | Amphotericin B-Norgestrel | | Norgestrel | Amphotericin B may decrease the excretion rate of Norgestrel which could result in a higher serum level. |
| HIV | Progesterone-Amphotericin B | | Progesterone | The metabolism of Amphotericin B can be decreased when combined with Progesterone. |
| HIV | Amphotericin B-Ethynodiol diacetate | | Ethynodiol diacetate | The metabolism of Amphotericin B can be increased when combined with Ethynodiol diacetate. |
| HIV | Diethylstilbestrol-Amphotericin B | | Diethylstilbestrol | The excretion of Amphotericin B can be decreased when combined with Diethylstilbestrol. |
| HIV | Amphotericin B-Ulipristal | | Ulipristal | The metabolism of Ulipristal can be decreased when combined with Amphotericin B. |
| HIV | Alitretinoin-Mifepristone | | Mifepristone | The therapeutic efficacy of Mifepristone can be decreased when used in combination with Alitretinoin. |
| HIV | Progesterone-Alitretinoin | | Progesterone | The metabolism of Alitretinoin can be decreased when combined with Progesterone. |
| HIV | Alitretinoin-Isoniazid | | Isoniazid | The therapeutic efficacy of Isoniazid can be decreased when used in combination with Alitretinoin. |
| Covid19 | Chloroquine-Heparin | | Heparin | The therapeutic efficacy of Chloroquine can be increased when used in combination with Heparin. |
| Covid19 | Methylprednisolone-Heparin | | Heparin | The therapeutic efficacy of Methylprednisolone can be increased when used in combination with Heparin. |
| Covid19 | Azithromycin-Fosinopril | | Azithromycin | The risk or severity of QTc prolongation can be increased when Fosinopril is combined with Azithromycin. |
| Covid19 | Azithromycin-Levobetaxolol | | Azithromycin | The risk or severity of QTc prolongation can be increased when Azithromycin is combined with Levobetaxolol. |
| Covid19 | Azithromycin-Practolol | | Azithromycin | The risk or severity of QTc prolongation can be increased when Azithromycin is combined with Practolol. |
| Covid19 | Azithromycin-Spirapril | | Azithromycin | The risk or severity of QTc prolongation can be increased when Spirapril is combined with Azithromycin. |
| Covid19 | Hyoscyamine-Ibuprofen | | Ibuprofen | The risk or severity of Tachycardia can be increased when Hyoscyamine is combined with Ibuprofen. |
| Covid19 | Pseudoephedrine-Ibuprofen | | Ibuprofen | The risk or severity of hypertension can be increased when Pseudoephedrine is combined with Ibuprofen. |
| Covid19 | Ibuprofen-Pimozide | | Ibuprofen | The risk or severity of hypertension can be increased when Ibuprofen is combined with Pimozide. |
| Covid19 | Ibuprofen-Hydrocortisone probutate | | Ibuprofen | The risk or severity of gastrointestinal irritation can be increased when Hydrocortisone probutate is combined with Ibuprofen. |
| Covid19 | Ibuprofen-Prednisolone acetate | | Ibuprofen | The risk or severity of gastrointestinal irritation can be increased when Prednisolone acetate is combined with Ibuprofen. |
| Covid19 | Dimenhydrinate-Ibuprofen | | Ibuprofen | The risk or severity of QTc prolongation can be increased when Dimenhydrinate is combined with Ibuprofen. |
| Covid19 | Ibuprofen-Albiglutide | | Ibuprofen | The risk or severity of angioedema can be increased when Ibuprofen is combined with Albiglutide. |
| Covid19 | Dexbrompheniramine-Ibuprofen | | Ibuprofen | The risk or severity of gastrointestinal bleeding can be increased when Ibuprofen is combined with Dexbrompheniramine. |
| Covid19 | Brompheniramine-Ibuprofen | | Ibuprofen | The risk or severity of gastrointestinal bleeding can be increased when Ibuprofen is combined with Brompheniramine. |
| Covid19 | Fosinopril-Dexamethasone | | Dexamethasone | The risk or severity of ulceration can be increased when Fosinopril is combined with Dexamethasone. |
| Covid19 | Sibutramine-Dexamethasone | | Dexamethasone | The risk or severity of myopathy and weakness can be increased when Dexamethasone is combined with Sibutramine. |
| Covid19 | Lisinopril-Dexamethasone | | Dexamethasone | The risk or severity of ulceration can be increased when Lisinopril is combined with Dexamethasone. |
| Covid19 | Temazepam-Dexamethasone | | Dexamethasone | The risk or severity of myopathy and weakness can be increased when Dexamethasone is combined with Temazepam. |
| Covid19 | Tamsulosin-Dexamethasone | | Dexamethasone | The risk or severity of myopathy, rhabdomyolysis, and myoglobinuria can be increased when Tamsulosin is combined with Dexamethasone. |
| Covid19 | Phentolamine-Dexamethasone | | Dexamethasone | The risk or severity of ulceration can be increased when Phentolamine is combined with Dexamethasone. |

**Supplemental Table 4. Predicted interactions between drugs used for women’s reproductive health and drugs used to treat HIV and COVID-19.**

Our model predicted fifteen and twenty-three interactions between drugs used for women’s reproductive health (WRHDs) and drugs approved to treat human immunodeficiency virus (HIV) and coronavirus disease 2019 (COVID-19), respectively. The predicted effect of each interaction is listed. DDI, drug-drug interaction.
